# Control of RNA polymerase II promoter-proximal pausing by DNA supercoiling

**DOI:** 10.1101/2020.05.12.091058

**Authors:** A. Herrero-Ruiz, P. Martínez-García, J. Terrón-Bautista, J.A. Lieberman, S. Jimeno-González, F. Cortés-Ledesma

**Author notes:** Lead Contact: Felipe Cortés-Ledesma.

## Abstract

The accumulation of topological stress in the form of DNA supercoiling is inherent to the advance of RNA polymerase II (Pol II) complexes, and needs to be resolved by DNA topoisomerases to sustain productive transcriptional elongation. Topoisomerases are therefore considered general positive facilitators of transcription. Here we show that, in contrast to this general assumption, human topoisomerase IIa accumulates at gene promoters, where it removes transcription-associated negative DNA supercoiling and represses transcription by enforcing promoter-proximal pausing of Pol II. We demonstrate that this topological balance is essential to maintain Immediate Early Genes under basal repression conditions, and that its disruption creates a positive feedback loop that explains their typical bursting behavior in response to stimulus. We therefore describe the control of promoter DNA supercoiling by topoisomerases as a novel layer for the regulation of gene expression, which can act as a molecular switch to rapidly activate transcription.

## Introduction

As originally envisioned by the ‘twin-supercoiled-domain’ model (Liu and Wang, 1987), the advance of RNA polymerase II (Pol II) during transcription is a torque-generating force that results in upstream and downstream regions of over- and underwound DNA, and thus positive (+) and negative (-) DNA supercoiling respectively (Kouzine et al., 2014; Ma and Wang, 2016). This supercoiling of template DNA can cause polymerase stalling (Ma et al., 2013), and therefore needs to be relieved in order to allow productive transcription and gene expression. DNA topoisomerases are the enzymes that relax this topological stress by transiently gating DNA passage, in a controlled cut-and-reseal mechanism that affects either one (type I DNA topoisomerases; mainly TOP1 in eukaryotes), or simultaneously both (type II topoisomerases; TOP2) DNA strands (Pommier et al., 2016). Hence, they are generally considered important facilitators of transcription, especially for long genes in which the load of DNA supercoiling can become particularly burdening (Joshi et al., 2012; King et al., 2013). In this sense, TOP1 and TOP2 have been shown to cooperate to relieve transcription associated (+) and (-) supercoiling in order to maintain appropriate levels of gene expression (Joshi et al., 2010; Pedersen et al., 2012; Sperling et al., 2011). It is worth noting, however, that DNA supercoiling can also facilitate transcription under some particular circumstances. For example, (-) supercoiling inherently helps duplex melting, the formation of Pol II open complexes and the recruitment of initiation factors (Parvin and Sharp, 1993; Tabuchi et al., 1993). Furthermore, the accumulation of (+) and (-) supercoiling has been proposed to facilitate the required nucleosome eviction ahead and reloading behind Pol II during transcription elongation (Corless and Gilbert, 2016; Teves and Henikoff, 2014). All this suggests that the regulation of supercoiling by topoisomerase activity could somehow operate to control transcription and gene expression, although the mechanisms by which this can occur remain largely unknown.

In mammals, TOP1 is essential for efficient transcription elongation and its activity is induced by direct interactions with the elongating form of Pol II (Baranello et al., 2016; Dujardin et al., 2014; King et al., 2013). In contrast, the functions of TOP2 in transcription seem more complex. Although also involved in facilitating elongation (King et al., 2013), both the a (TOP2A) and β (TOP2B) mammalian TOP2 paralogs are enriched at promoters (Canela et al., 2017; Thakurela et al., 2013; Tiwari et al., 2012), suggesting relevant regulatory functions. Indeed, TOP2A and TOP2B control expression of neuronal differentiation genes that are essential for proper neural development (Lyu et al., 2006; Thakurela et al., 2013; Tiwari et al., 2012). Accumulating evidence also suggests that TOP2B can be essential for the fast induction of highly regulated genes in response to different types of stimuli, including hormones (Haffner et al., 2010; Ju et al., 2006) growth factors (Bunch et al., 2015) and neuronal activity (Madabhushi et al., 2015). Immediate Early Genes (IEG) such as *c-FOS*, which respond with a transient burst in transcription only a few minutes after cells are stimulated, are a paradigm of this type of transcriptional response (Healy et al., 2013). Interestingly, this has been proposed to operate through the generation of a TOP2-mediated DSB at promoters, which would be directly responsible for locally triggering transcription (Bunch et al., 2015; Ju et al., 2006; Madabhushi et al., 2015). As mentioned above, TOP2 enzymes indeed cut duplex DNA as part of their catalytic cycle, remaining covalently linked to the ends of the incised fragment in the so-called cleavage complex (TOP2cc) (Pommier et al., 2016). These intermediates, however, are very rare in cells, unless stabilized with TOP2 poisons such as etoposide (Canela et al., 2017; Gittens et al., 2019; Gothe et al., 2019), and are fully reversible structures that only result in DSB formation upon interference with cellular processes and proteasomal degradation (Canela et al., 2019; Gothe et al., 2019; Sciascia et al., 2020; Zhang et al., 2006). In addition, there is profuse evidence for DSBs, both in promoter and gene bodies, as being strong inhibitors of transcription *in cis*, in a process that depends on DNA-damage response signaling, facilitates repair of the lesions and protects genome integrity (Caron et al., 2019; Hanawalt and Spivak, 2008; Iacovoni et al., 2010; Pankotai et al., 2012; Shanbhag et al., 2010). The TOP2-DSB model of transcription stimulation is therefore difficult to reconcile with classic knowledge on TOP2 biology and the DNA-damage response.

Alternative hypotheses to explain the function of TOP2 and DNA supercoiling in the control of gene expression should therefore be considered. Although in many cases gene regulation implies changes in the recruitment of Pol II to promoters and increased transcription initiation, in mammals, the main regulatory level lies at the entry into productive transcription elongation (Adelman and Lis, 2012). This is exerted through the modulation of promoter-proximal pausing, a transient arrest that Pol II suffers shortly after initiating RNA synthesis, remaining primed for a transition into elongation upon P-TEFb-mediated Ser2 phosphorylation of its C-terminal domain (CTD) (Adelman and Lis, 2012; Core and Adelman, 2019; Li and Gilmour, 2011). For this reason, global analyses of Pol II occupancy show that a large fraction of genes display Pol II signal concentrated near the TSS (Guenther et al., 2007; Kim et al., 2005). Promoter-proximal pausing is controlled by specific complexes such as Negative Elongation Factor (NELF) and DRB sensitivity inducing factor (DSIF) that decrease elongation efficiency of Pol II (Yamaguchi et al., 2013), as well as by intrinsic features of the template chromatin (Aoi et al., 2020; Jimeno-Gonzalez et al., 2015; Kwak et al., 2013). Interestingly, promoter-proximal pausing has been recently shown to correlate with TOP2B and spontaneously occurring DSBs at a genome-wide level (Dellino et al., 2019), and interpreted in the context of the TOP2B-mediated DSB model of transcriptional regulation.

Here we unexpectedly find that the main effect of TOP2 catalytic inhibition in human cells is a quick and global release of Pol II from promoter-proximal pausing that results in a sharp upregulation of immediate early and other highly regulated genes. Interestingly, this is a result of TOP2A repressive functions in transcriptional regulation that are tightly interconnected to TOP1 activity and completely independent of DSB formation, but instead, rely on the continuous removal of transcription-associated (-) supercoiling at promoter regions. With these results, we provide a comprehensive topological framework for the regulation of promoter-proximal pausing that explains the typical bursting behavior of IEGs, and in more general terms, the tight control of human gene expression by DNA topoisomerases.

## Results

### TOP2 catalytic inhibitors induce expression of IEGs

To study a possible function of TOP2 and DNA supercoiling in regulating transcription, changes in gene expression profiles were analyzed in human telomerase-immortalized retinal pigment epithelial cells (RPE-1) treated with merbarone, a drug that catalytically inhibits TOP2 upstream of DNA cleavage (Fortune and Osheroff, 1998). Cells were previously arrested in G0/G1 by serum starvation in order to avoid possible effects of topoisomerase inhibition on other cellular processes such as replication or chromosome segregation. Upon a 2-hour treatment, the comparison between RNA-seq profiles of merbarone- and mock-treated samples revealed that 173 protein-coding genes were differentially expressed, with 148 upregulated and only 25 downregulated upon merbarone treatment (Figure 1A; Data S1), pointing to a mainly repressive role of TOP2 activity on gene expression. Interestingly, most upregulated genes (UP genes) were related with cellular responses to different cellular stimuli (Figure S1A). In fact, there was a clear enrichment of IEGs (Figure S1B), genes that, as mentioned above, are highly regulated, and activated through protein synthesis-independent rapid bursts of transcription shortly after different types of stimuli (Bahrami and Drablos, 2016). A paradigm of this pattern of expression is the *c-FOS* gene, whose mRNA production peaks at 30-60 min after stimulation and drops after 90 min (Greenberg and Ziff, 1984). Therefore, we analyzed RNA-seq samples taken 30 minutes after merbarone treatment and found that only 5 genes were upregulated: *c-FOS, FOSB, ATF3, ZFP36* and *JUNB*; all of them well-established IEGs (Figure 1B; Data S1). The transcriptional de-repression of *c-FOS*, which we will use as a representative model from now on, and *EGR1*, another responsive IEG, under TOP2 inhibition was confirmed by reverse transcription quantitative PCR (RT-qPCR), with an outstanding induction (80- and 10-fold, respectively) 30 min after merbarone treatment and a subsequent decrease at later time points (Figure 1C) that mirrored physiological induction with serum (Figure S1C). This behavior was not particular to RPE-1 cells, being also observed in the lung carcinoma A549 cell line and in primary Mouse Embryonic Fibroblasts (MEFs) (Figure S1D and S1E), demonstrating conservation among cell types and species. Importantly, RPE-1 cells treated with ICRF-187, a different TOP2 catalytic inhibitor that targets TOP2 after religation, also showed *c-FOS* upregulation (Figure 1D). In contrast, treatment with the paradigmatic TOP2 poison etoposide resulted in a minor induction of *c-FOS* expression only at later time points (Figure 1E), similar to that previously reported in other cell types (Bunch et al., 2015; Madabhushi et al., 2015), but neglible when compared to that observed upon treatment with merbarone or physiological induction with serum. We can therefore conclude that treatment with TOP2 catalytic inhibitors results in a rapid and robust induction of highly regulated genes, including a subset of IEGs, and *c-FOS* in particular.

**Figure 1.**
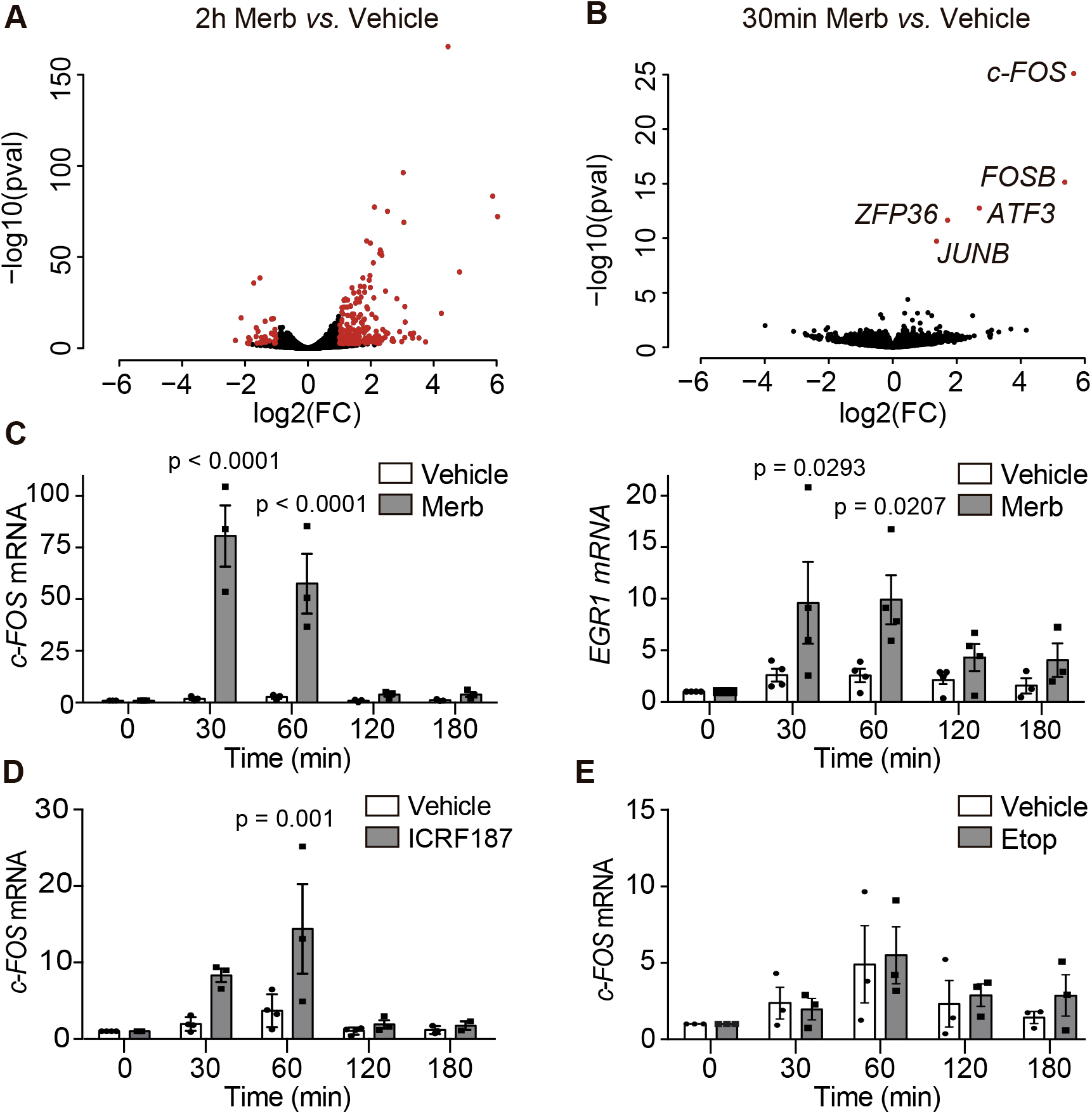
Immediate Early Genes (IEGs) are upregulated upon TOP2 catalytic inhibition. (A and B) Volcano plots of transcriptional changes upon 2 hours (A), or 30 minutes (B), merbarone (Merb, 200 μM) versus vehicle (DMSO) treatments measured by RNA-seq on serum-starved RPE-1 cells. The x axis represents fold-induction ratios in a log-2 scale and y axis represents p-value a log-10 scale. Genes with an absolute fold change ≥2 and an adjusted p-value ≤ 0.05 are shown in red. (C) *c-FOS* (left) and *EGR1* (right) mRNA levels, as measured by RT-qPCR, at the indicated times following merbarone and vehicle (DMSO) treatments (*n=3*). Values were normalized to *GAPDH* mRNA levels and signal in untreated conditions. Individual experimental values and mean ± s.e.m. are shown, two-way ANOVA tests with Bonferroni post-test. (D) As in (C) for *c-FOS* expression upon ICRF-187 (200 μM) treatment. (E) As in (C) for *c-FOS* expression upon.etoposide (Etop, 20 μM) treatment.

### IEG induction is independent of DSBs or stress

The results described above strongly suggest TOP2 inhibition as a potential molecular mechanism for IEG upregulation, challenging current models of TOP2-mediated promoter DSBs (Bunch et al., 2015; Madabhushi et al., 2015). There is, however, some degree of controversy as to whether merbarone can also act as a TOP2 poison *in vivo* (Pastor et al., 2012). We therefore decided to analyze the accumulation of TOP2ccs and DSBs under conditions of *c-FOS* upregulation. First, we performed ICE (*In vivo* Complex of Enzyme), which measures TOP2ccs by isolating and detecting covalent protein-DNA complexes accumulated in cells (Nitiss et al., 2012) (Figure S2A). Clearly, neither merbarone-nor serum-treatments resulted in detectable accumulation of TOP2ccs (Figure 2A). Second, we monitored the generation of DSBs by the quantification of gH2AX foci by immunofluorescence (Kinner et al., 2008). Again, we found that neither merbarone nor serum produced a significant accumulation of gH2AX foci (Figure 2B), despite concomitantly causing a robust transcriptional upregulation of *c-FOS* (Figure 2C). In contrast, treatment with etoposide resulted in the expected strong induction of both TOP2ccs and DSBs without a significant induction of *c-FOS* transcription (Figure 2A-C). The transcriptional response, therefore, does not correlate with the appearance of TOP2ccs and DSBs. To further confirm that IEG induction was independent of DNA damage, we decided to impair the repair of TOP2-associated DSBs by removing TDP2, a highly specialized DNA-end unblocking enzyme (Cortes Ledesma et al., 2009) whose absence significantly delays repair of this type of DNA lesion (Gomez-Herreros et al., 2013; Schellenberg et al., 2017). As a matter of fact, RPE-1 cells deleted for *TDP2* by CRISPR-Cas9 displayed a kinetics of *c-FOS* induction indistinguishable from wild-type cells, both upon merbarone and serum treatments (Figure S2B and S2C). Altogether, these results strongly disfavor the involvement of stable TOP2ccs or associated DSBs in the mechanism of transcriptional upregulation of IEGs. Interestingly, this is not only true for merbarone treatment, but also for physiological stimulation with serum.

**Figure 2.**
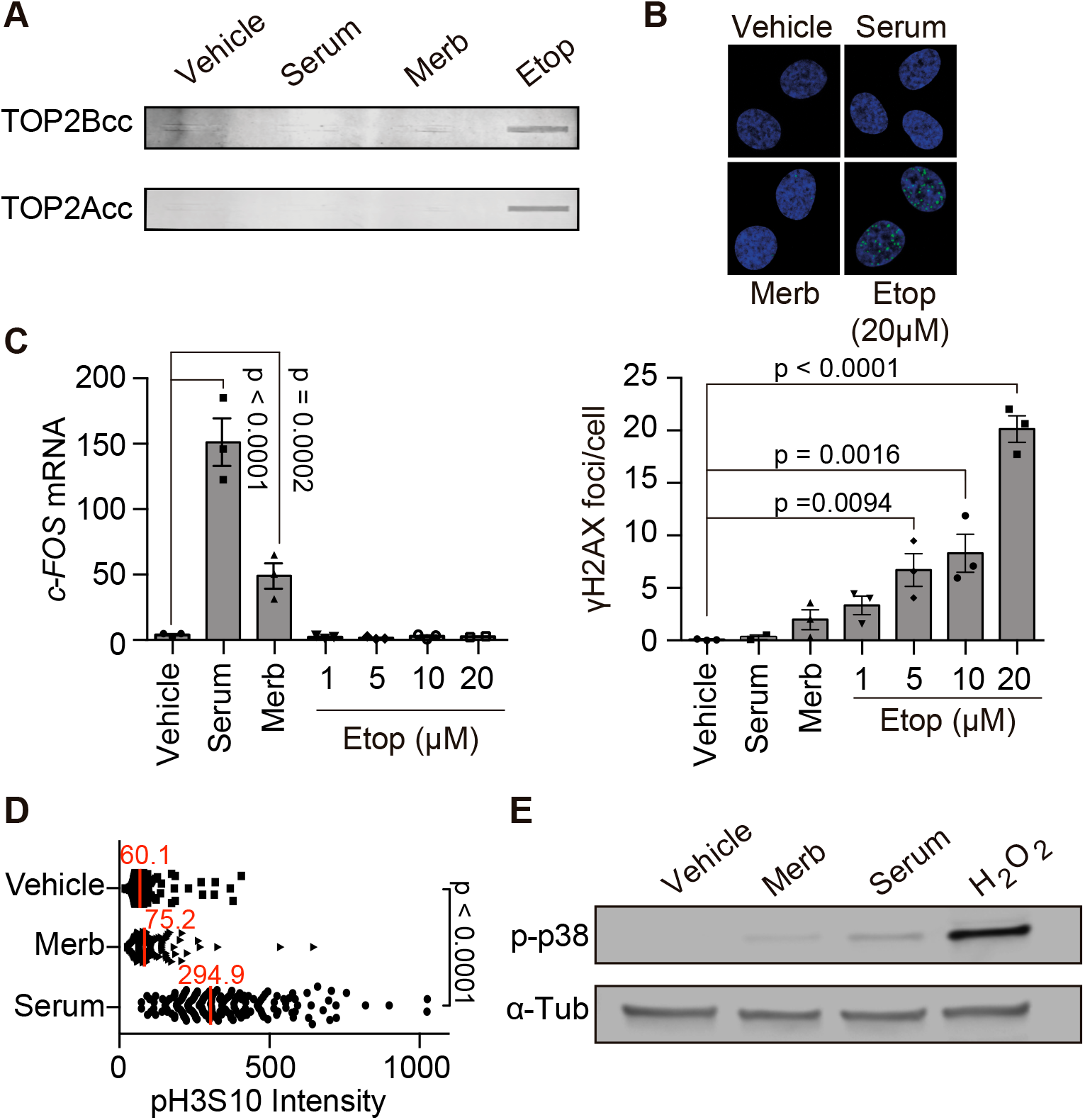
IEG induction is independent of DSBs or stress. (A) TOP2A and TOP2B covalently bound to genomic DNA (ICE) under 30 minutes of vehicle (DMSO), merbarone (Merb, 200 μM), serum (1%) or etoposide (Etop, 20 μM) treatments. A representative image is shown (*n=3*). (B) Representative image (top) and quantification (bottom) of γH2AX foci (green) immunofluorescence upon 30 minutes of the indicated conditions (*n=3*). DAPI counterstaining is shown (blue). Individual experimental values and mean ± s.e.m. are shown, one-way ANOVA tests with Bonferroni post-test. (C) *c-FOS* mRNA levels upon 30 minutes of the indicated conditions (*n=3*), details as in Figure 1C. (D) Quantification (down) of phosphorylated H3S10 (pH3S10) immunofluorescence in serum-starved RPE-1 cells treated for 1 h with vehicle (DMSO), merbarone (Merb, 200 μM) or serum (1%) (*n=3*). Individual cellular values and mean ± s.e.m., two-way ANOVA tests with Bonferroni post-test. (E) Western blot of p-p38 MAPK (Thr180/Thr182) upon 30 minutes incubation with vehicle (DMSO), merbarone (Merb, 200 μM), serum (1%) or H2O2 (2 μM). A representative image is shown (*n=3*).

Finally, we decided to test whether the transcriptional response to merbarone could be an indirect cause of some type of cellular stress or signaling derived from TOP2 inhibition. There are two main pathways responsible for IEG upregulation in response to stimulus: the RAS-MAPK pathway that is activated by growth factors and mitogens, and the p38-MAPK pathway that is activated by UV or other types of stress (Healy et al., 2013). Strikingly, and in contrast to physiological induction with serum or hydrogen peroxide, we found that merbarone treatment did not significantly increase phosphorylation of histone H3 serine 10 (H3S10) (Figure 2D and S2D) or p38 (Figure 2E), which are well-established makers of the respective activation of these pathways. Hence, we conclude that merbarone increases IEG expression directly, bypassing activation of canonical signaling pathways, and not by indirectly eliciting a cellular stress response, although we cannot fully rule out the involvement of additional yet-unknown signaling pathways.

### TOP2A activity at gene promoters represses transcription of IEGs

To characterize the inhibitory effect of merbarone on TOP2 activity, ICE experiments were carried out following different times of merbarone treatment and a brief (5 min) incubation with high dose (400 μM) etoposide in order to “freeze” catalytically engaged enzymes (Figure 3A). In this experimental setup, etoposide-mediated TOP2cc induction is used as an estimate of TOP2 activity, which merbarone is expected reduce by acting upstream in the catalytic cycle. Strikingly, at times in which IEG upregulation was already evident, merbarone mainly inhibited the TOP2A paralog, while TOP2B activity was only mildly affected at later time points (Figure 3B). Consistent with this, stable in-pool deletion of *TOP2B* in RPE-1 cells overexpressing Cas9 (RPE-1 Cas9) (Figure S3A) did not change *c-FOS* expression when compared to cells transfected with a non-targeting gRNA, neither its basal levels (Figure 3C), nor its induction upon merbarone or serum treatments (Figure 3D). Importantly, these result not only prove that TOP2B is not a target for the transcriptional response elicited by merbarone, but also rule out the previously proposed requirement for TOP2B-mediated DSBs in the regulation of *c-FOS* expression (Bunch et al., 2015; Madabhushi et al., 2015), at least in our experimental conditions. In order to check the involvement of TOP2A, an essential protein for cell cycle progression, we performed acute *TOP2A* deletion in conditions of serum starvation by the transfection of an appropriate gRNA in RPE-1 Cas9 cells (Figure S3B). Despite these technical limitations, we achieved substantial levels (>90%) of TOP2A protein depletion (Figure 3E), which were sufficient to cause a significant increase in the basal levels of *c-FOS* when compared to cells transfected with a non-targeting gRNA (Figure 3E). This was evident even in these particular experimental conditions, which already resulted in some degree of constitutively increased *c-FOS* expression (Figure S3C), possibly due to the stress of cellular transfection. In any case, treatment with both merbarone or serum were still capable of triggering substantial *c-FOS* upregulation in control cells (Figure 3F). In *TOP2A*-deleted cells, however, merbarone-mediated *c-FOS* induction was completely abolished, while stimulation with serum was potently enhanced (Figure 3F). These results strongly support inhibition of TOP2A as the direct cause of *c-FOS* upregulation in response to merbarone treatment, ruling out potential off-target effects of the drug. Furthermore, the de-repression observed in *TOP2A*^-/-^ cells, both in terms of basal expression levels and upon serum stimulation, suggests that merbarone mainly operates through a reduction in TOP2A activity and not by indirect effects caused by the trapping of the enzyme on chromatin.

**Figure 3.**
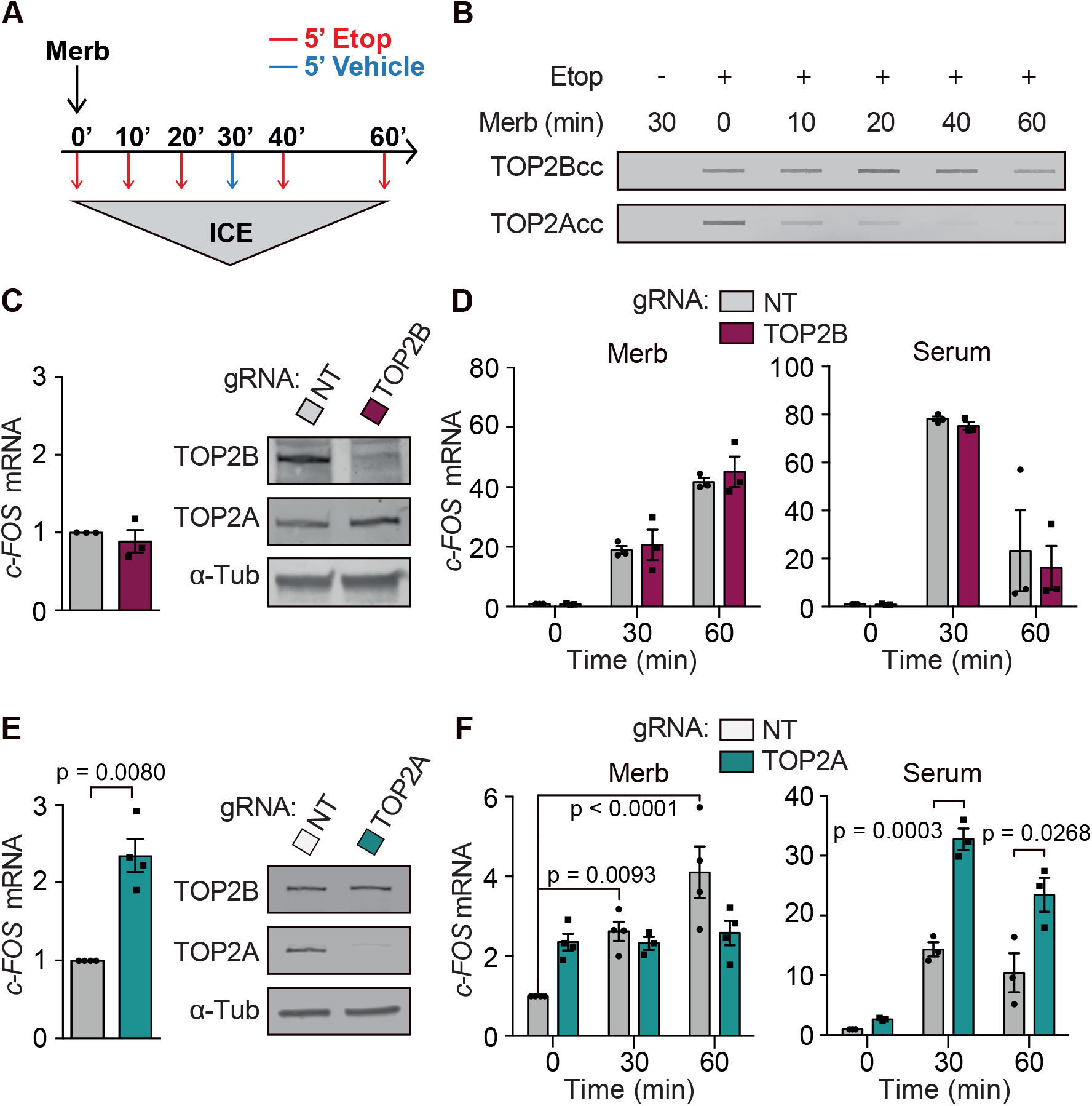
Merbarone induces IEGs by inhibiting TOP2A. (A, B) Diagram of the experimental design (A), and a representative image (B) of ICE experiment to measure the inhibitory effect of merbarone (Merb, 200 μM) on TOP2 catalytic activity (*n=3*). 48h serum-starved (0.1%) RPE-1 cells were subjected to merbarone treatment for the indicated time with a final 5 minutes treatment with either vehicle (DMSO) or etoposide (400 μM). (C) *c-FOS* mRNA levels in RPE-1 Cas9 cells following transfection with non-targeting (NT) and *TOP2B-specific* (TOP2B) gRNAs (*n=3*), according to the experimental outline illustrated in Figure S3A. Individual values and mean ± s.e.m. are shown, One-sample t-test (hypothetic value = 1). A representative western-blot image of TOP2B and TOP2A levels in WT and *TOP2B*^-/-^ RPE-1 cells is shown (right) (*n=3*). (D) *c-FOS* mRNA levels in the cells described in (C) following the indicated times of merbarone (left) or serum (right) treatments (*n=3*), details as in Figure 1C. (E) As in (C) for a *TOP2A*-specific (TOP2A) gRNAs (*n=3*). Experimental outline in Figure S3B (F) As in (D) for a *TOP2A*-specific (TOP2A) gRNAs (*n=3*).

Direct transcriptional roles of TOP2A are somewhat unexpected, as these have been traditionally assigned to TOP2B (Austin et al., 2018). However, TOP2A has been reported to physically interact with Pol II (Mondal and Parvin, 2001), and to accumulate at promoters and nucleosome-free regions (Canela et al., 2017; Thakurela et al., 2013), similarly to TOP2B. ChIP-seq in G0/G1-arrested RPE-1 cells confirmed this enrichment at promoters and enhancers in our experimental conditions (Figure 4A). In fact, the distribution of TOP2A strongly correlated with Pol II and marks of active chromatin like H3K4me3 and H3K27ac at these regions (Figure S4). Interestingly, average TOP2A levels around the transcription-start site (TSS) were notably higher in the 148 genes upregulated at the mRNA level after 2h merbarone treatment (UP genes) than in the same number of randomly selected genes (Figure 4B). This is consistent with a direct involvement of TOP2A in transcriptional repression of these genes under basal conditions. TOP2A profiles in *c-FOS* and *EGR1*, as positive examples, and *LDLR*, an IEG not upregulated upon merbarone treatment (Data S1), illustrate these differences (Figure 4C). To further link TOP2A activity to transcriptional repression, we decided to determine whether TOP2A present at the *c-FOS* promoter was catalytically active. To do so, we set up a technique (TOP2A ICE-IP) in which accumulated TOP2ccs were immunoprecipitated from ICE extracts with specific antibodies against TOP2A, and the associated DNA was subsequently amplified by qPCR with different pairs of primers along the *c-FOS* gene (Figure S2A and Figure 4D). Interestingly, etoposide treatment strongly increased the amount of covalently bound TOP2A at the *c-FOS* promoter, indicative of its strong catalytic engagement at this region. TOP2A is therefore, not only present but active at the *c-FOS* promoter under basal conditions, in agreement with a repressive role in the transcriptional regulation of this gene. We therefore conclude that TOP2A plays a major role in the basal constitutive repression of *c-FOS*, which is overcome upon inhibition of its catalytic activity, leading to a sharp transcriptional upregulation.

**Figure 4.**
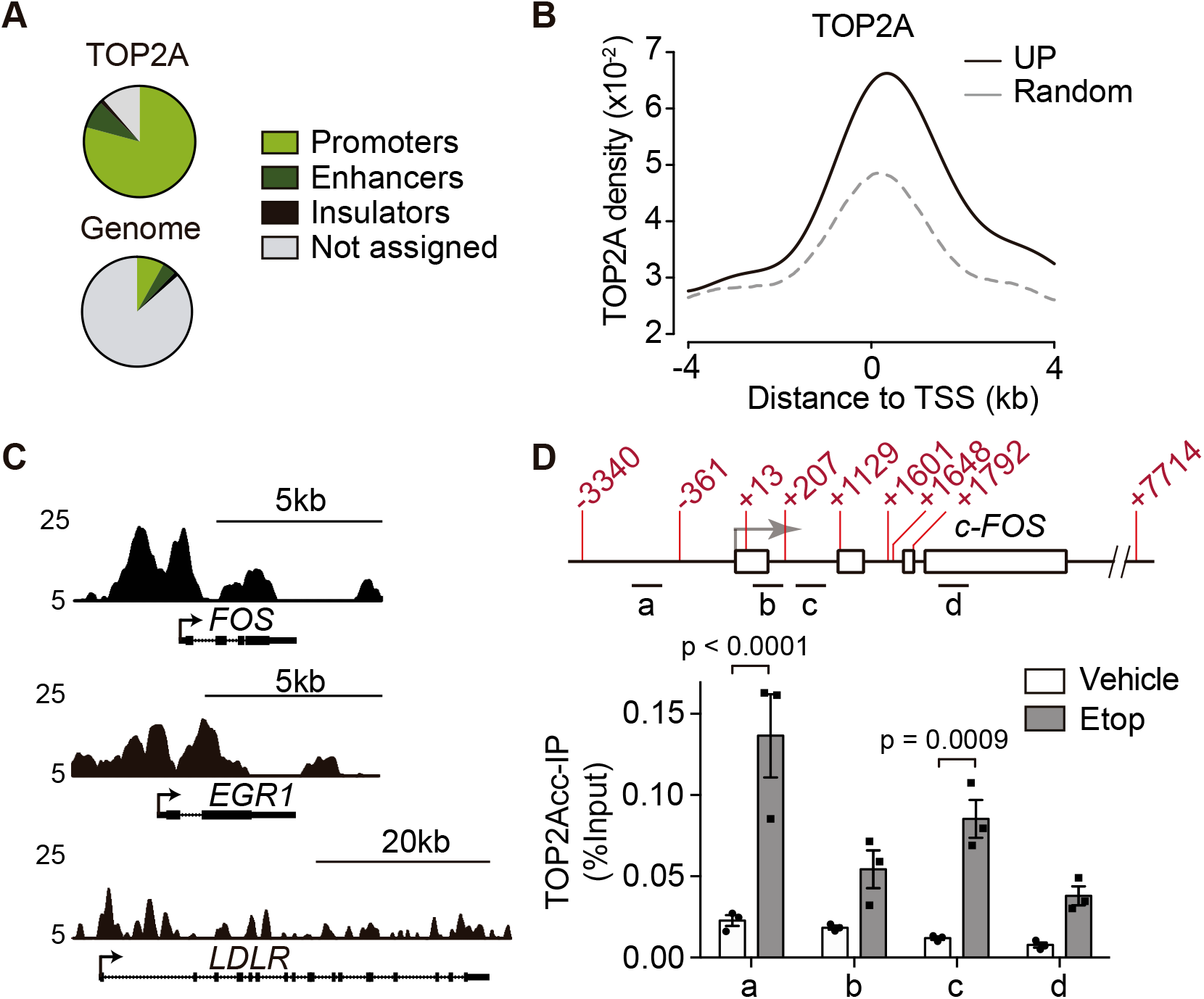
TOP2A activity at gene promoters represses transcription. (A) Genome-wide distribution of TOP2A peaks compared to the general distribution of human mappable genomic regions. (B) Average TOP2A ChIP-seq enrichment around the TSS of UP genes and an equal number of randomly selected genes. (C) Genome browser tracks for TOP2A ChIP-seq at *c-FOS, EGR1* and *LDLR* genes in untreated serum-starved RPE-1 cells. The y axis represents RPKM. (D) TOP2A ICE-IP under vehicle (DMSO) or etoposide (Etop) (400 μM) treatment for 5 minutes. A scheme of the amplified regions corresponding to Pst I fragments in the *c-FOS* gene (top), and TOP2Acc relative enrichment (bottom) (*n=3*), are shown. Data are individual experimental values and mean ± s.e.m., two-way ANOVA tests with Bonferroni post-test.

### TOP2A catalytic inhibition releases Pol ll from promoter-proximal pausing

In order to understand the transcriptional effect of TOP2A inhibition in detail, we used ChIP-seq to compare the distribution of Pol II in control conditions and following 30 and 60 minutes of merbarone treatment. An analysis of differential Pol II gene body occupancy showed a repression of only 4 genes, while 48 were induced at 60 minutes of merbarone treatment, with 19 of them (18%) coinciding with the genes upregulated at the mRNA level (UP genes) (Figure S5A and S5B). In fact, Pol II occupancy at the body of UP genes was significantly increased upon merbarone treatment (Figure 5A), confirming that the effect of TOP2A inhibition on gene expression was mainly exerted at the level of transcriptional upregulation. This effect could be clearly observed in *c-FOS* and *EGR1* as representative examples, but not in the irresponsive *LDLR* IEG (Figure 5B). Strikingly however, while Pol II was significantly increased in the gene body of UP genes, this was not the case in the region surrounding the TSS, where it was apparently decreased, although not reaching statistical significance (Figure 5A). This phenotype is consistent with transcriptional upregulation being mainly caused by an increased release from promoter-proximal pausing, as can be observed in the profile of Pol II occupancy (Figure 5C). The decrease at the TSS was, however, not observed in strongly induced genes such as *c-FOS* and *EGR1* (Figure 5B), probably because of a concomitant stimulation of transcription initiation (Shao and Zeitlinger, 2017). To quantify the changes in Pol II distribution, two gene body/promoter ratios were calculated, Pause Release Ratio (PRR) (Chen et al., 2015) and Pausing Index (PI) (Core et al., 2008; Day et al., 2016), which are positive and negative indicators of pause release, respectively. Merbarone treatment significantly increased PRR (Figure 5C) and decreased PI (Figure S5C) of UP genes, suggesting that TOP2A inhibition leads to gene upregulation through the release of Pol II from the pause site. We then decided to extend our analysis to all (n=10,471) active genes, as determined by the presence of Pol II at the promoter in control conditions. Again, although more mildly, average Pol II gene occupancy showed a general shift from the TSS to proximal coding regions, and significant increases in PRR (Figure 5D) and decreases in PI (Figure S5D) under TOP2A inhibition. Finally, we also evaluated the genome-wide effect of merbarone treatment on promoter-proximal pausing using an empirical cumulative distribution function (ECDF) of PRR, which resulted in an evident shift towards pause release (Figure 5E). We therefore conclude that TOP2A inhibition leads to an increase in the release of Pol II from promoter-proximal pausing that can be observed globally, although only a subset of genes becomes significantly upregulated at the mRNA level.

**Figure 5.**
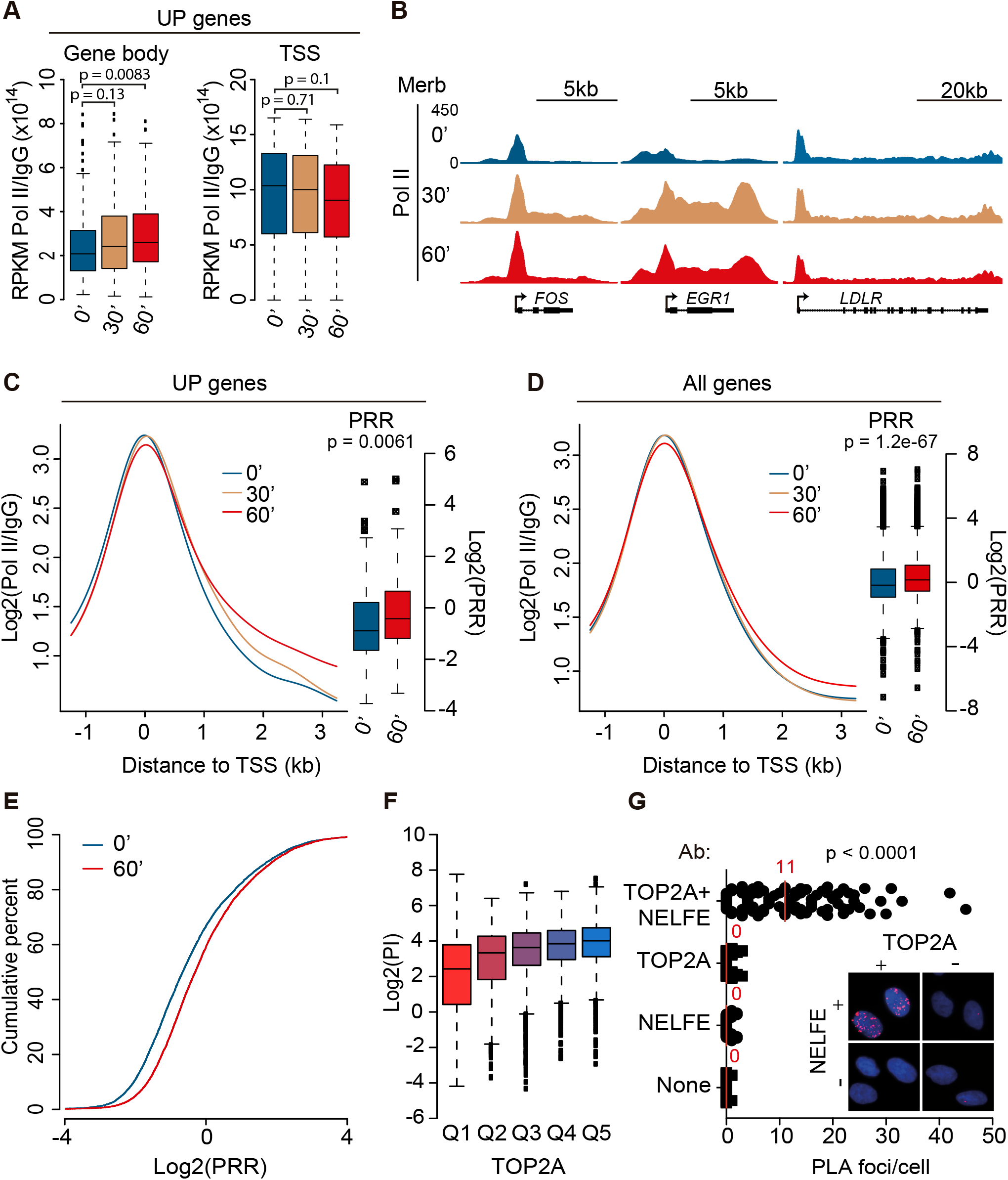
Merbarone releases Pol ll from promoter-proximal pausing. (A) Box-plot distribution of Pol II/IgG ChIP-seq reads (RPKM) at the gene body (+0.5 to +2.5 kb; left) and the TSS (−0.5 to +0.5; right) in UP genes following the indicated times of merbarone treatment. Wilcoxon rank-sum test (B) Genome browser tracks for Pol II ChIP-seq (top) following the indicated times of merbarone treatment at *c-FOS, EGR1* and *LDLR* genes. The y axis represents RPKM. (C) Average Pol II ChIP-seq distribution around the TSS (left) and box-plot distribution of PRR (right) in UP genes following the indicated times of merbarone treatment. Wilcoxon rank-sum test. (D) As in (C) for all (10,471) transcriptionally active genes, determined by Pol II or H3K4me3 ChIP-seq signal at the TSS. (E) Empirical cumulative distribution function (ECDF) of PRR following the indicated times of merbarone treatment. (F) Correlation between TOP2A levels at the TSS and PI. Genes were stratified in quintiles (Q1 to Q5) regarding their TOP2A levels at the TSS, as measured by TOP2A ChIP-seq counts, and the box-plot distribution of PI in each quintile is represented. (G) Proximity ligation assay (PLA) quantification and representative image (inset) of TOP2A and NELFE interaction in U2OS cells. The antibody used in each condition is indicated on the y axis. Individual cells and median is shown (40 cells; n=3), one-way ANOVA test with Bonferroni post-test.

To further link TOP2A to the regulation of promoter-proximal pausing, genes were categorized in quintiles regarding their TOP2A density at promoters and their average PI and PRR were determined. Interestingly, TOP2A positively correlated with PI (Figure 5F) and negatively with PRR (Figure S5E), indicating a direct association of TOP2A with promoter-proximal pausing. Furthermore, we found that TOP2A was in close proximity to NELF, as determined by proximity-ligation assay (PLA) against its RNA-binding subunit NELFE (Figure 5G). The specificity of this signal was confirmed by the reduction of PLA foci under NELFE downregulation (Figure S5F). These results, together with the general pause release and transcriptional upregulation effects observed under merbarone treatment, strongly suggest that TOP2A is an important repressor of transcription that operates by a stimulation of promoter-proximal pausing.

### *c-FOS* expression is regulated by DNA supercoiling

TOP2 and TOP1 have been shown to display redundant functions in the removal of transcription-associated supercoiling, at least in yeast cells (Joshi et al., 2012; Pedersen et al., 2012; Sperling et al., 2011). We therefore decided to monitor changes in TOP1 activity during *c-FOS* upregulation, by using TOP1 ICE-IP, as described above for TOP2A but with a pulse of the TOP1 poison camptothecin (CPT) (Figure S2A). We found that TOP1 had a basal activity that was highly increased within the *c-FOS* coding region when transcription was induced with serum (Figure 6A), consistent with the reported tight link between TOP1 activity and transcription elongation (Baranello et al., 2016). Upon merbarone treatment, however, TOP1 activity in *c-FOS* remained at basal levels (Figure 6A), despite the dramatic increase in transcriptional activity (Figure 1C). Importantly, this was not due to off-target effects of merbarone on TOP1 function, as global CPT-induced TOP1cc levels were not altered (Figure S6A). Thus, TOP2A inhibition completely bypasses the need of TOP1 for transcription elongation, a result that confirms the topological nature of merbarone-induced gene upregulation. Conversely, TOP1 inhibition with CPT completely suppressed *c-FOS* upregulation upon merbarone treatment (Figure 6B). We can conclude that, TOP2A and TOP1, rather than operating redundantly, are negative and positive transcriptional effectors, respectively, and whose equilibrium is essential to regulate *c-FOS* expression. Anyhow, the fact that an acute transcriptional response can be achieved without an accompanying increase in topoisomerase activity is indeed remarkable.

**Figure 6.**
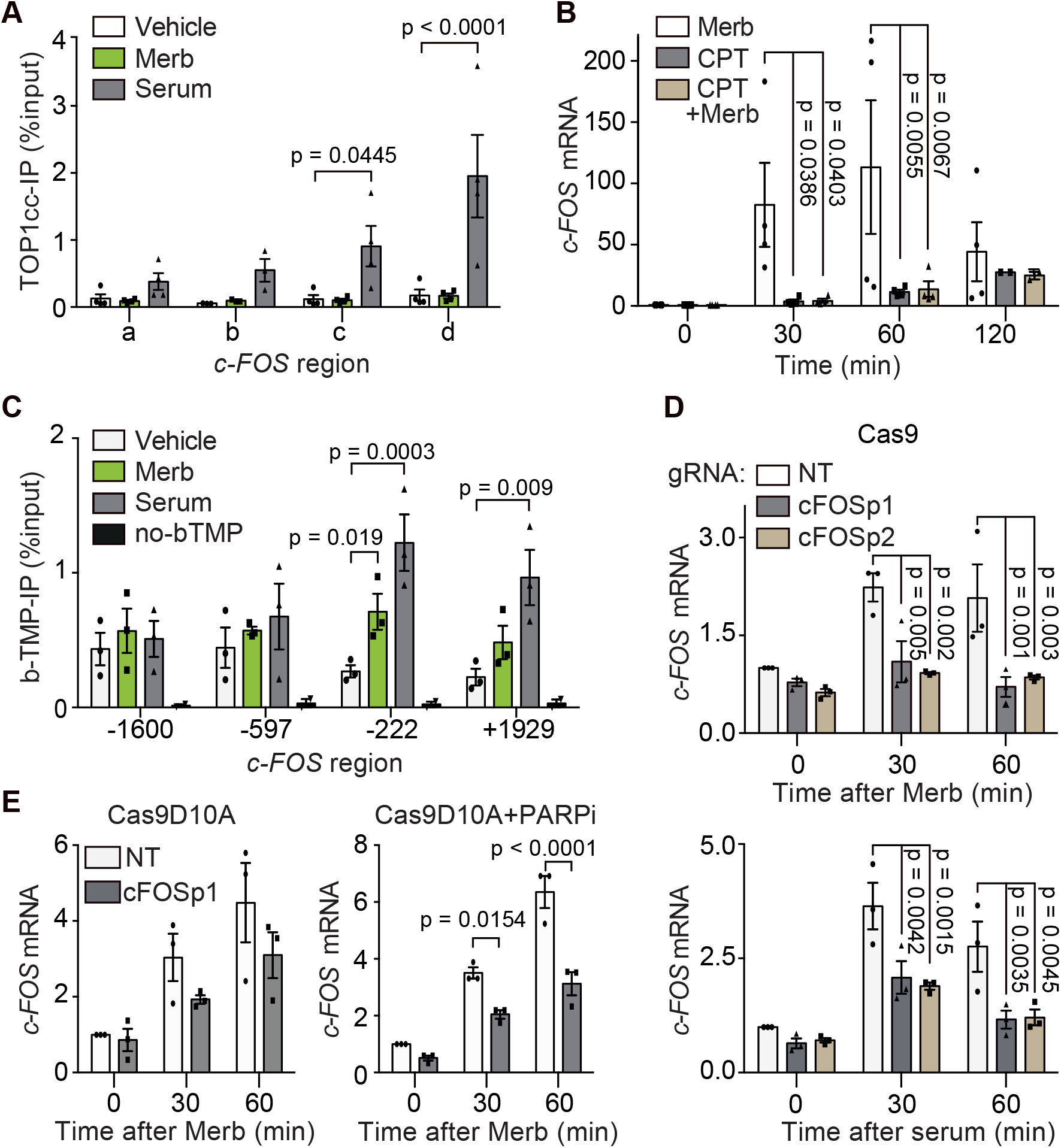
c-FOS expression is regulated by DNA supercoiling. (A) TOP1cc levels, as determined by TOP1 ICE-IP, at the indicated regions of the *c-FOS* gene (see Figure 4D) following 30 minutes vehicle (DMSO), merbarone (Merb, 200μM) or serum (1%) treatments. In all conditions, TOP1ccs were induced by 5 minutes incubation with camptothecin (CPT, 10 μM) (*n=4*). Individual experimental values and mean ± s.e.m. are shown, two-way ANOVA test with Bonferroni post-test. (B) *c-FOS* mRNA levels in RPE-1 cells upon treatment with merbarone (Merb, 200μM), camptothecin (CPT, 10 μM) or their combination, as indicated (*n=3*), details as in Figure 1C. (C) Biotin-trimethyl-psoralen (bTMP) incorporation, expressed as relative enrichment after pull-down, at the indicated positions of the *c-FOS* gene in serum-starved RPE-1 cells after 10 min treatments with vehicle (DMSO), merbarone (Merb, 200μM) or serum (1%) (*n=3*). Untreated cells without bTMP incubation (no-bTMP) are included as a negative control. Individual experimental values and mean ± s.e.m. are shown, two-way ANOVA test with Bonferroni post-test. (D) *c-FOS* mRNA levels in RPE-1 Cas9 cells 2 h after transfection with the indicated gRNA, and at different times following induction with merbarone (Merb, 200μM) (top) or serum (1%) (bottom) (*n=3*), other details as in Figure 1C. (E) As in (D) but in RPE-1 Cas9 D10A cells, and including 30 min pretreatment with PARP1 inhibitor PJ34 (PARPi;10 μM; right).

The involvement of two opposing topoisomerase activities strongly suggests a central role for DNA topology, and likely DNA supercoiling, in the regulation of *c-FOS* expression. We therefore decided to monitor changes in the incorporation of biotinylated trimethyl psoralen (bTMP) (Figure S4A), a compound that preferentially intercalates into negatively-supercoiled (-) DNA (Bermudez et al., 2010), during conditions of *c-FOS* upregulation (Figure S6B). Indeed, stimulation of *c-FOS* transcription by merbarone treatment resulted in an increased incorporation of bTMP at the *c-FOS* promoter, similarly to what was observed upon physiological induction with serum (Figure 6C). This suggests that (-) supercoiling accumulates at the *c-FOS* promoter region during transcriptional stimulation, and is consistent with genome-wide observations in yeast and mammalian systems (Bermudez et al., 2010; Kouzine et al., 2013; Naughton et al., 2013). However, we considered the possibility that this accumulation of (-) supercoiling could not only represent a consequence of transcription, but play an active role in its regulation. In order to tackle this question, we decided to disrupt local supercoiling by the induction of DNA breaks. We individually transfected two different gRNAs in serum-starved RPE-1 Cas9 cells to target a DSB to the *c-FOS* promoter region (Figure S6C), and analyzed changes in gene expression with merbarone and serum. Interestingly, in both cases *c-FOS* induction was strongly disrupted upon transfection with the targeting gRNAs, and not when a non-targeting control was used (Figure 6D). These results further argue against the DSB model of transcriptional stimulation, and suggest, instead, that the topological integrity of the promoter region is an essential factor to allow gene expression. In addition, since DSBs are known to strongly repress transcription *in cis* (Caron et al., 2019), we decided to alternatively target a CRISPR-Cas9 D10A nickase that generates single-strand breaks (SSBs) (Chiang et al., 2016). Again, we observed a disruption of merbarone-mediated *c-FOS* induction, which was further enhanced and reached statistical significance when signaling and repair were impaired upon treatment with PARP1 inhibitor PJ34 (Figure 6E). We conclude that the accumulation of (-) supercoiling at the promoter region is necessary for *c-FOS* transcriptional stimulation. Furthermore, altogether, the results confirm the disruption of DNA topology as the molecular cause of merbarone-induced transcription upregulation, and suggest a tight interdependence between TOP2A and TOP1 activities in order to maintain supercoiling homeostasis and transcriptional control.

## Discussion

### A model of supercoiling-mediated regulation of promoter-proximal pausing

The results presented here demonstrate that the continuous removal of (-) supercoiling from gene promoters by TOP2A facilitates promoter-proximal pausing of Pol II, and that this is essential to maintain *c-FOS* and other IEGs under basal repressed conditions. It may seem counter-intuitive that a topoisomerase, which relieves torsional stress, acts to negatively regulate transcription. One must bear in mind however that (-) supercoiling is a well-established stimulator of transcription initiation, at least *in vitro* and in prokaryotic models (Chong et al., 2014; Kim et al., 2019; Parvin and Sharp, 1993; Revyakin et al., 2004; Tabuchi et al., 1993), and has been proposed to operate similarly in yeast and mammalian cells (Baranello et al., 2016; Bermudez et al., 2010). Thus, (-) supercoiling at promoter regions facilitates transcription initiation by allowing promoter melting and the formation of Pol II open complexes. In this same line, although upstream (-) supercoiling can stall bacterial RNA polymerase (Ma et al., 2013), when encountered downstream, it can favor transcriptional elongation by cancelling the (+) supercoiling generated (Kim et al., 2019). In this context, we propose that, by removing (-) supercoiling generated behind advancing Pol II complexes, TOP2A limits initiation and processivity of subsequent transcriptional cycles, imposing a strong requirement for TOP1-mediated removal of (+) supercoiling ahead of each elongating polymerase (Figure 7). This is particularly relevant in the vicinity of the TSS, where TOP1 activity is reduced (Baranello et al., 2016) (Figure 6A), favoring the formation of a (+) topological barrier that enforces promoter-proximal pausing and maintains transcription under repressed and controlled conditions. Disrupting this balance, either in a regulated manner or by inhibiting TOP2A activity, leads to a transcription-dependent accumulation of (-) supercoiling, in a topological feedback loop (transcription increases supercoiling and supercoiling increases transcription) that overcomes promoter-proximal pausing and can result in strong transcriptional bursts, particularly in genes, such as IEGs, that are particularly poised for transcriptional stimulation, and in which pausing is the limiting step. In fact, the equilibrium between Top I and gyrase activities in removing (-) and (+) supercoils, respectively, has been shown to regulate transcriptional bursting in bacteria (Chong et al., 2014).

**Figure 7.**
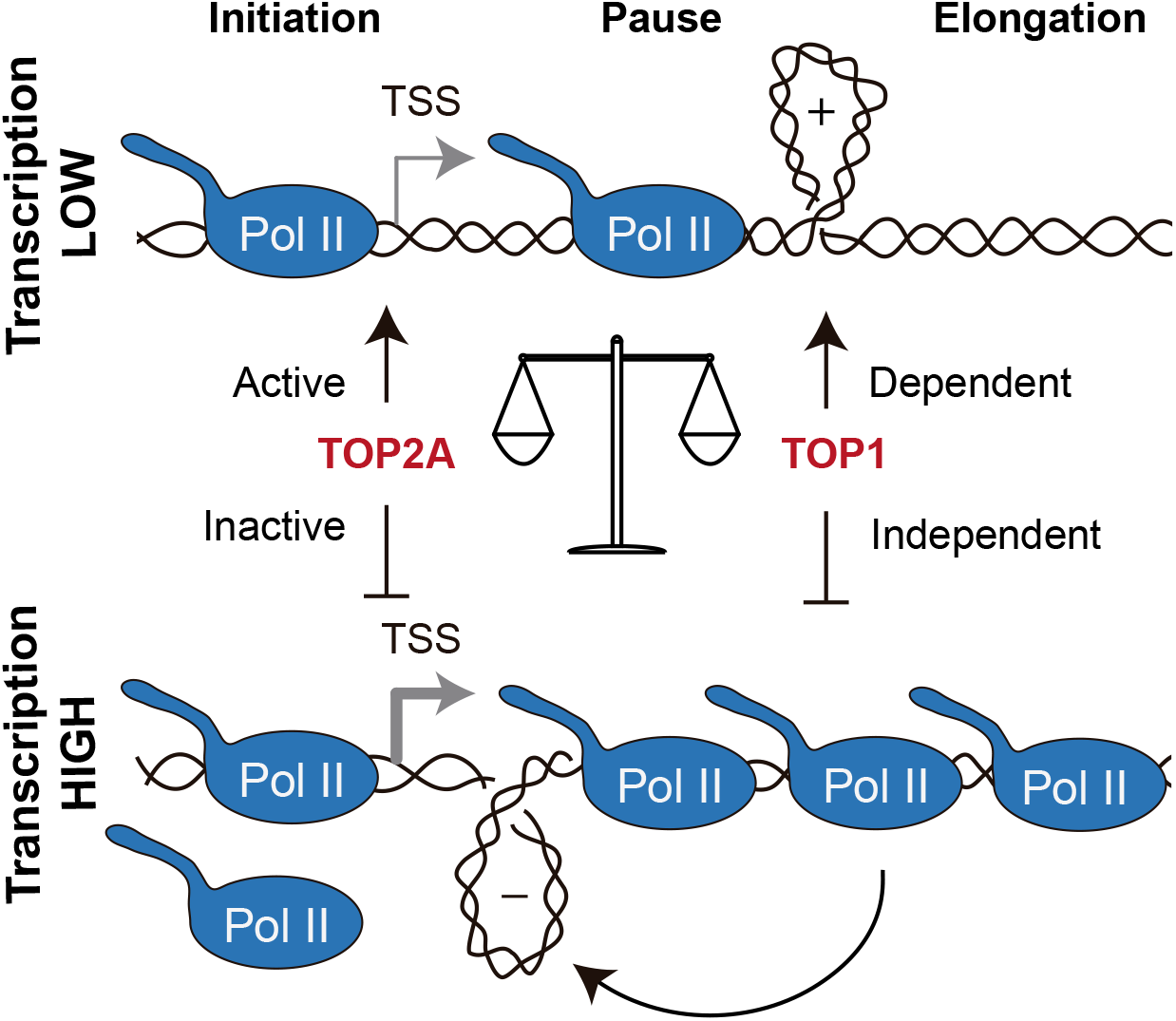
A model of supercoiling-mediated regulation of promoter proximal pausing. Proposed model for transcription regulation through DNA supercoiling. Under repressive conditions, Pol II early elongation generates (+) DNA supercoiling ahead and (-) DNA supercoiling behind. TOP1 removes (+) DNA supercoiling to favour Pol II pause release, but the removal of promoter (-) DNA supercoiling by TOP2A maintains transcription under controlled conditions. If TOP2A is inactive, (-) DNA supercoiling accumulated at the promoter counteracts (+) DNA supercoiling, allowing TOP1-independent advance of Pol II, and bypassing promotor-proximal pausing.

It is worth noting that in these conditions of absent or limited TOP2A-mediated (-) supercoiling removal, multiple Pol II complexes could advance in tandem, with a cancellation of (+) and (-) supercoiling between them, in “trains” of polymerases in which only the “engine” encounters topological constraints, so upregulation of transcription can be achieved by simply adding more “wagons” without additional topological costs. This could explain how merbarone treatment leads to a strong transcriptional stimulation of *c-FOS* without increased TOP1 activity (Figure 6A). Interestingly, long-range communication between RNA polymerases by supercoiling cancelation and interference has been recently reported in bacteria (Kim et al., 2019). In any case, the accumulation of (-) supercoiling at promoter regions seems a wide-spread characteristic of eukaryotic gene organization (Bermudez et al., 2010; Kouzine et al., 2013; Matsumoto and Hirose, 2004; Naughton et al., 2013; Teves and Henikoff, 2014), and could serve additional functions such as, for example, favoring the formation of non-B DNA structures with regulatory functions (Kouzine et al., 2004; Kouzine et al., 2008), providing an appropriate 3D conformation to favor interaction with regulatory elements (Liu et al., 2001; Racko et al., 2019) or as a means to isolate and maintain the topology of the transcriptional unit (Achar et al., 2020). Finally, the tight interdependence between chromatin structure, DNA supercoiling and topoisomerase function (Dykhuizen et al., 2013; Kaczmarczyk et al., 2020; Sperling et al., 2011; Teves and Henikoff, 2014) provides an additional level of complexity expanding the possibilities of fine-tune transcriptional regulation.

It is perhaps shocking that, despite a general release from promoter-proximal pausing upon merbarone treatment, similar to that caused by depletion of established pausing factors (Chen et al., 2015; Core et al., 2012), only a relatively small number of genes is upregulated (Figure 1A). One should bear in mind, however, that pausing of Pol II has been demonstrated to be a necessary step for optimal transcription. In fact, upon depletion of promoter-proximal pausing factors, transcriptional upregulation is not widespread, but mainly restricted to rapidly inducible genes, like heat-shock or, interestingly, immediate-early genes (Fujita et al., 2009; Gilchrist et al., 2008; Schaukowitch et al., 2014). In this sense, promoter-proximal pausing also indirectly regulates Pol II recruitment to promoters by inhibiting transcription re-initiation (Shao and Zeitlinger, 2017). This might be especially relevant for those highly-regulated genes that are characterized by fast and synchronous changes in gene expression under stimulation, in which not only elongation but also initiation has to be rapidly increased. TOP2A inhibition may therefore operate in a similar fashion, with expression levels being significantly altered only in genes, such as IEGs, that are poised for transcription stimulation and lack other repressive mechanisms of regulation. Furthermore, this suggests that DNA supercoiling is one of many aspects contributing to promoter-proximal pausing of Pol II, which may thus be regulated at different levels. The relative contribution of these factors could explain the variable response of individual genes, cell types or tissues.

### TOP2-mediated DSBs and transcription regulation: cause or consequence?

Our model radically changes current view of IEG stimulation through the physiological induction of TOP2B-mediated DSBs at gene promoters (Bunch et al., 2015; Madabhushi et al., 2015), an idea that was originally put forward for the regulation of hormone responsive genes (Ju et al., 2006). In contrast, we provide an alternative possibility by which IEG expression is stimulated by (-) DNA supercoiling, and actually repressed by the action of TOP2A. Indeed, we present compelling evidence demonstrating that neither TOP2A-nor TOP2B-mediated DSBs are necessary for IEG expression. First, we are unable to detect TOP2ccs or DSBs upon merbarone or serum stimulation, despite a concomitant strong induction of transcription. Second, we show that *c-FOS* stimulation does not change in cells deleted for the highly TOP2-specialized repair enzyme TDP2. Although alternative TDP2-independent pathways to repair TOP2-induced DSBs exist in cells, they operate at a slower rate and seriously compromise genome integrity (Alvarez-Quilon et al., 2014; Gomez-Herreros et al., 2013), and would therefore inevitably impact on the extent and/or kinetics of putative DSB-dependent gene expression. Third, neither TOP2A nor TOP2B deletion reduce *c-FOS* expression or stimulation with serum, demonstrating the complete dispensability of their activity. As a matter of fact, TOP2A-deleted cells display higher basal *c-FOS* expression and a more accused serum response, in agreement with its repressive functions. Finally, we show that DSBs targeted to the *c-FOS* promoter, rather than stimulating, abolish its capacity to respond to merbarone or serum treatments.

Conciliatingly however, although it is difficult for us to fully explain some of the results that support the TOP2-DSB model, some key observations are actually completely compatible. Thus, a mild (~2-fold) induction of *c-FOS* and other IEGs was found at late times of etoposide treatment in neurons (Madabhushi et al., 2015). Indeed, here we observe a very similar behavior in RPE-1 cells (Figure 1E), but the fact that a much higher and faster induction is observed upon catalytic inhibition (Figures 1C and 1D), strongly suggests the reduction in activity, which also occurs upoon etoposide treatment, rather the generation of TOP2ccs or DSBs, as the molecular trigger of the transcriptional response. Furthermore, the association between transcription and DNA damage is clear and well documented (Gaillard and Aguilera, 2016), and it is plausible that at least part of this can be caused by accidental topoisomerase-mediated DNA breaks (Sun et al., 2020). We therefore favor a scenario in which TOP2-induced DSBs are accidental, and a consequence, rather than the cause, of transcriptional upregulation. In support for this, chemical inhibition of transcription elongation dramatically reduces the accumulation of spontaneous DSBs that normally colocalize with TOP2B and Pol II at gene promoters (Dellino et al., 2019).

## Conclusions

Our results uncover a new layer of transcriptional regulation that operates at the level of promoter-proximal pausing and depends on promoter supercoiling regulated by the balance between TOP2A and TOP1 activities. In addition, our discoveries provide a molecular explanation for the typical bursting behavior of immediate early, and potentially other highly regulated genes. Interestingly, this topological control operates at a high hierarchical level, being sufficient to trigger gene expression overcoming other regulatory steps, such as signaling cascades, chromatin remodeling or the recruitment of specific factors. Furthermore, our findings place the functions of TOP2 in transcriptional regulation in a context, that, in contrast to previous models based on DSB formation (Ju et al., 2006; Madabhushi et al., 2015), are aligned with classical knowledge on TOP2 catalysis, and importantly, do not unnecessarily compromise genome integrity. In more general terms, this study constitutes a first step to understand how TOP2 activity, and DNA supercoiling in general, may be used to modulate biological processes in mammalian cells. Future work should be devoted to study the regulation of topoisomerase function and DNA supercoiling in this and other relevant physiological scenarios, and how they converge with different aspects of chromatin dynamics to shape genome function, organization and stability.

## Acknowledgements

We thank the Genomics Core facilities at CABIMER and the EMBL (GeneCore) for the generation of the high-thoughput sequencing data and O. Fernández-Capetillo for comments. Computational analyses were run on the High Perfomance Computing cluster provided by the Centro Informático Científico de Andalucía (CICA). This work was funded with grants from the Spanish and Andalusian Governments (SAF2017-89619-R, CVI-7948, European Regional Development Fund), and the European Research Council (ERC-CoG-2014-647359); and with individual fellowships for A.H-R. (Contratos para la Formación de Doctores, BES-2015-071672, Ministerio de Economía y Competitividad), S.J-G. (Ramón y Cajal, RYC-2015-17246, Ministerio de Economía y Competitividad) and J.T-B. (Formación Profesorado Universitario, FPU15/03656, Ministerio de Educación, Cultura y Deporte). CABIMER is supported by the Andalusian Government.

## Author Contributions

A.H-R., S.J-G. and F.C-L. conceived and designed the project. A.H-R., P.M-G., S.J-G. and J.T-B performed the experiments and analyzed the results. S.J-G. and F.C-L. wrote the manuscript. All authors read, discussed and approved the manuscript.

## Figure Legends

**Figure S1.**
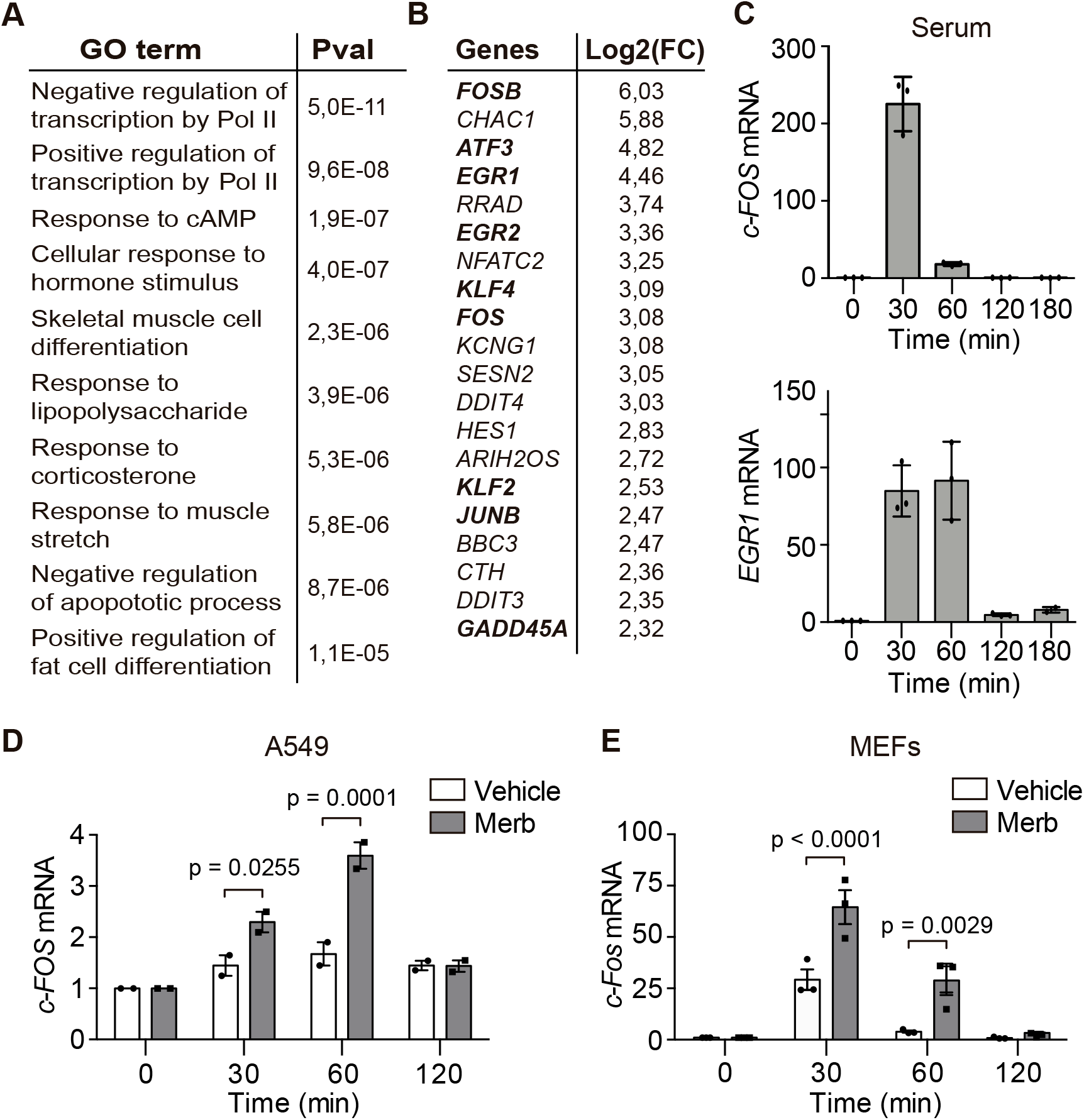
Changes in gene expression upon merbarone treatment and serum in different cell types. (A) Top 10 significantly enriched Gene Ontology (GO) biological process terms for UP genes. (B) Top 20 UP genes sorted by fold change. Early-Response Genes are in bold. (C) *c-FOS* (top) and *EGR1* (bottom) mRNA levels in serum-starved RPE-1 cells at the indicated times after serum (1%) addition (n=3), details as in Figure 1C. (D) *c-FOS* mRNA levels in serum-starved A549 cells at the indicated times after merbarone (Merb, 200 μM) or vehicle (DMSO) treatments (n=3), details as in Figure 1C. (E) As in (D) for *c-Fos* mRNA in Mouse Embryonic Fibroblasts (MEFs)

**Figure S2.**
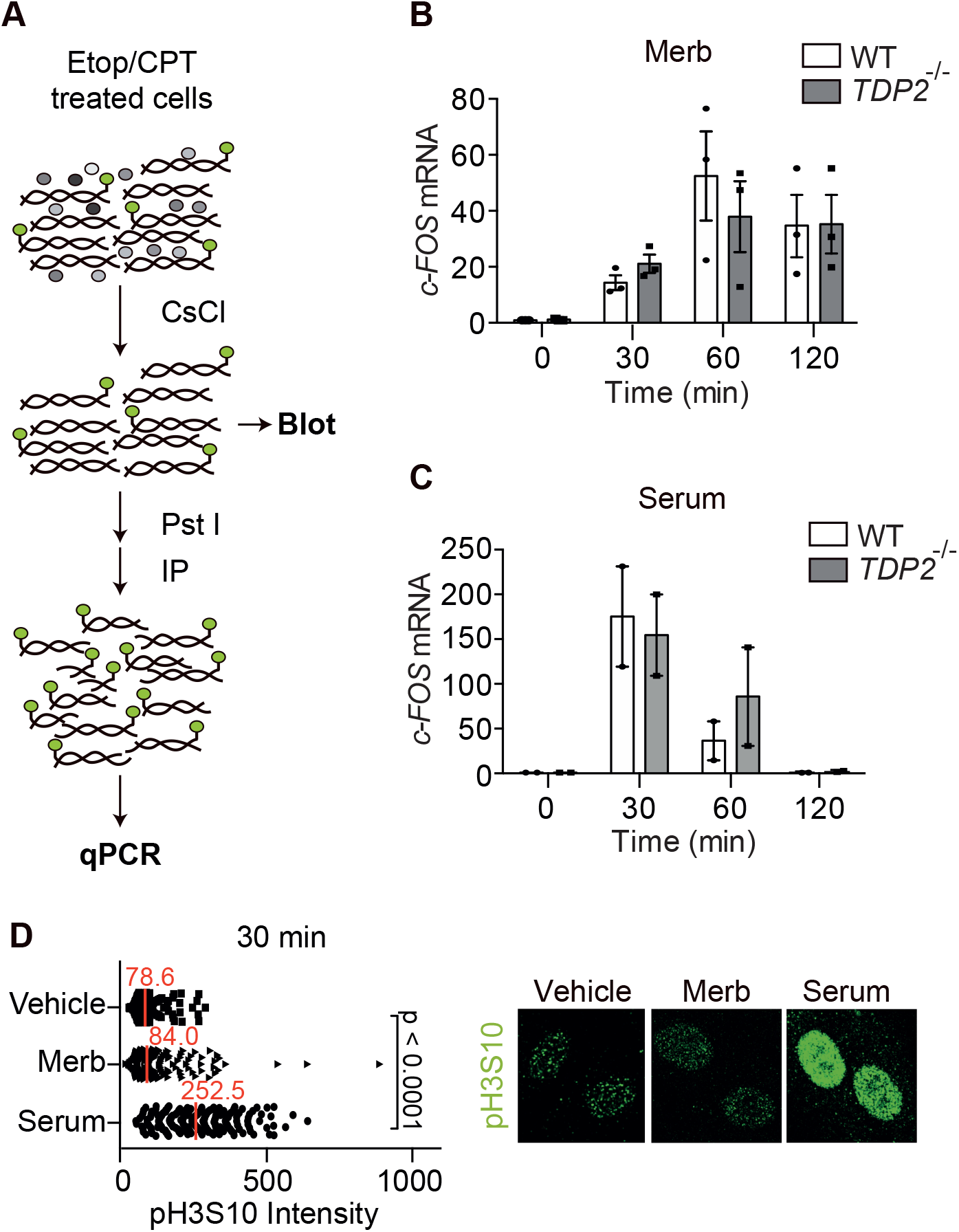
Analysis of TOP2cc, DSBs and stress response. (A) Schematic representation of *In vivo* Complex of Enzyme (ICE) and ICE-seq experimental set-up. (B and C) *c-FOS* mRNA levels in WT and *TDP2*^-/-^ serum-starved RPE-1 cells at the indicated times after the addition of merbarone (n=3) (B) or serum (n=2) (C), other details as in Figure 1C. (D) Quantification (left) and representative imagen (right) of phosphorylated H3S10 (pH3S10) immunofluorescence in serum-starved RPE-1 cells treated for 30 min with vehicle (DMSO), merbarone (Merb, 200 μM) or serum (1%) (*n=3*), details as in Figure 2D.

**Figure S3.**
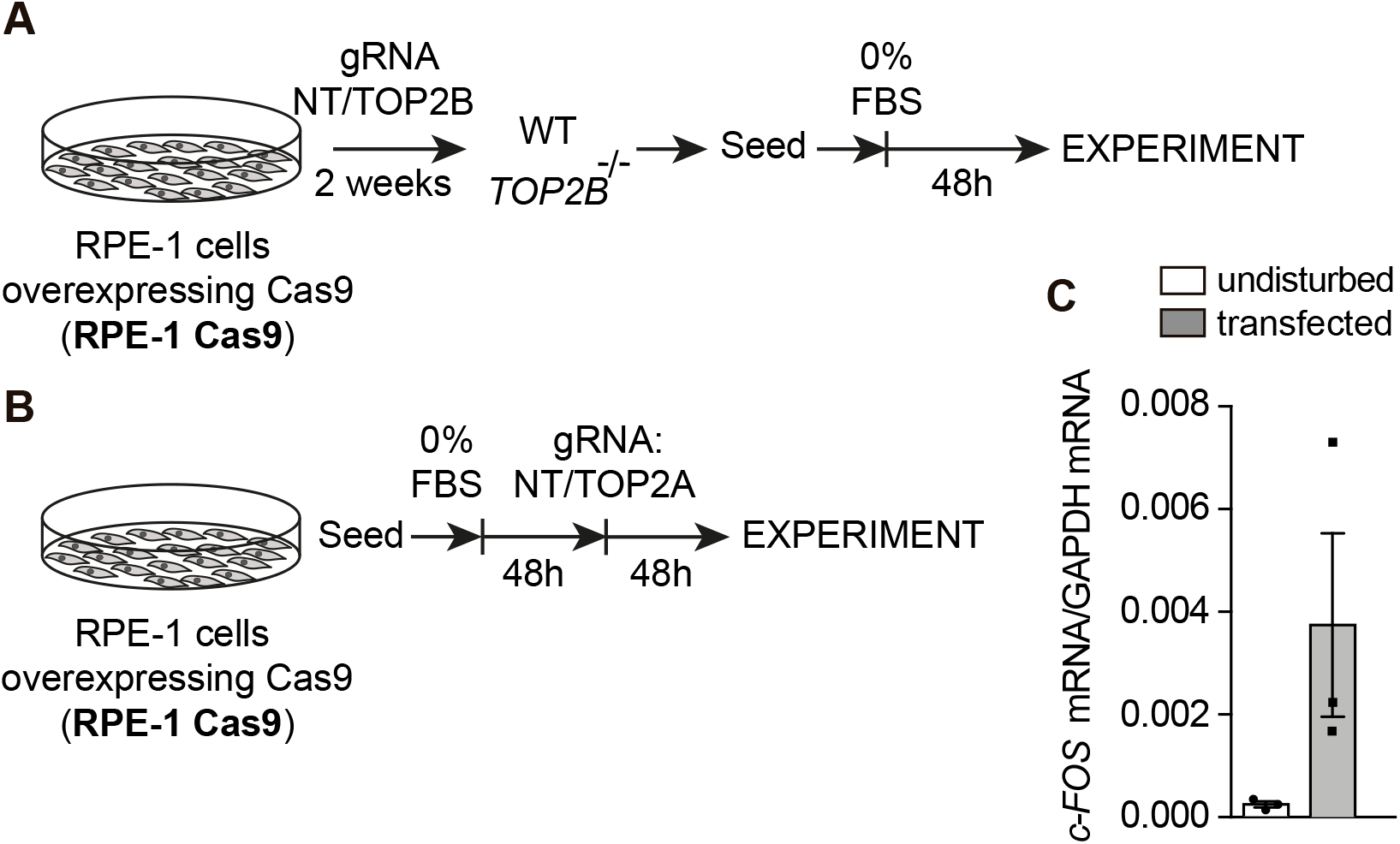
CRISPR-Cas9 deletion of TOP2 paralogs. (A) Scheme of strategy for the generation of *TOP2B*^-/-^ RPE-1 cells. (B) Scheme of strategy for the generation of *TOP2A*^-/-^ RPE-1 cells. (C) Absolute c-*FOS* mRNA/GAPDH mRNA levels in undisturbed cells and 48 h after non-targeting gRNA transfection (*n=3*). Individual experimental values and mean ± s.e.m. are shown.

**Figure S4.**
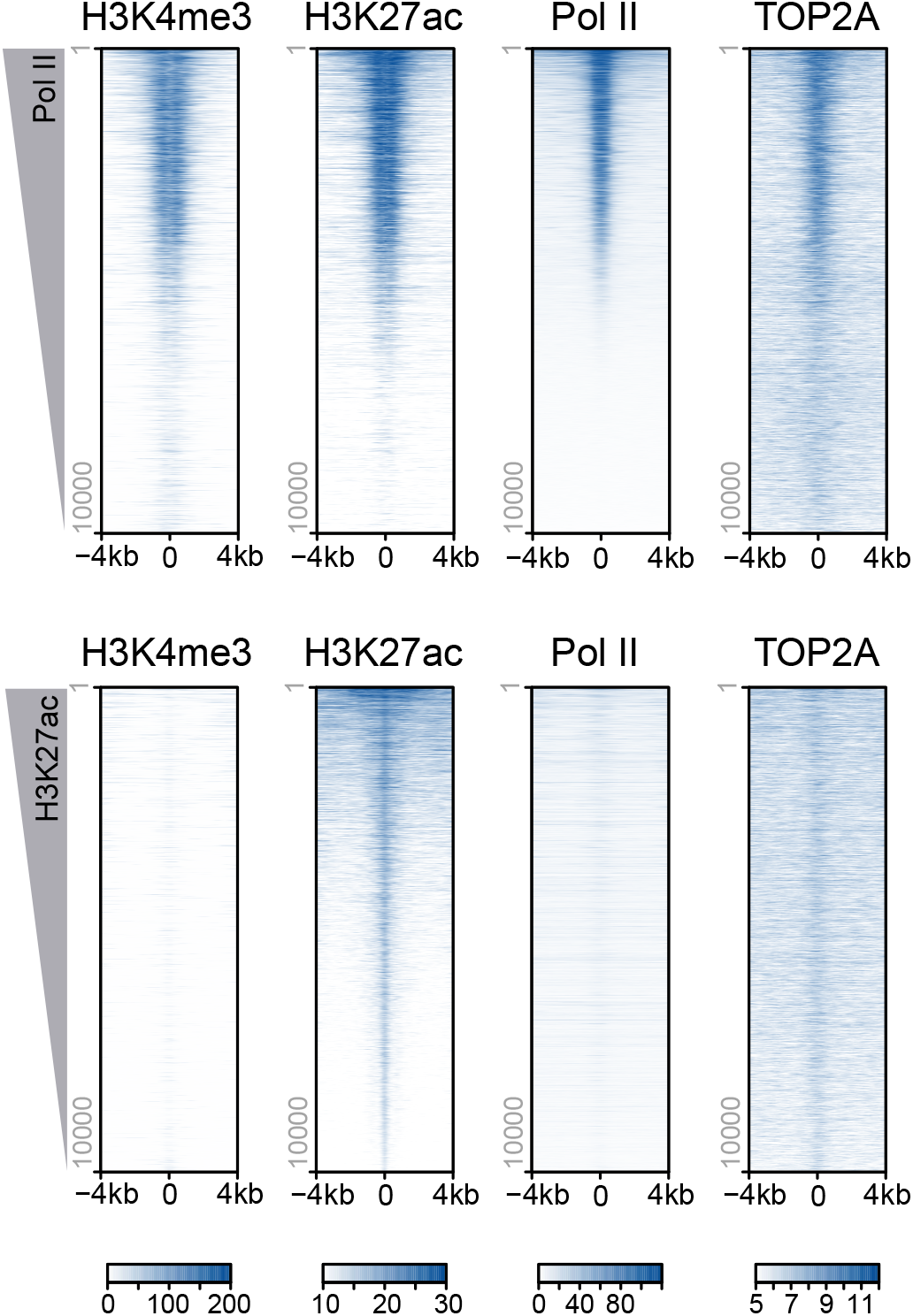
TOP2A is enriched at active chromatin regions. Heatmaps displaying H3K4me3, H3K27ac and TOP2A ChIP-seq signal in a ± 4kb region centered on promoters (top) and enhancers (bottom). Decreasing Pol II and H3K27ac signals are used to order promoters and enhancers, respectively.

**Figure S5.**
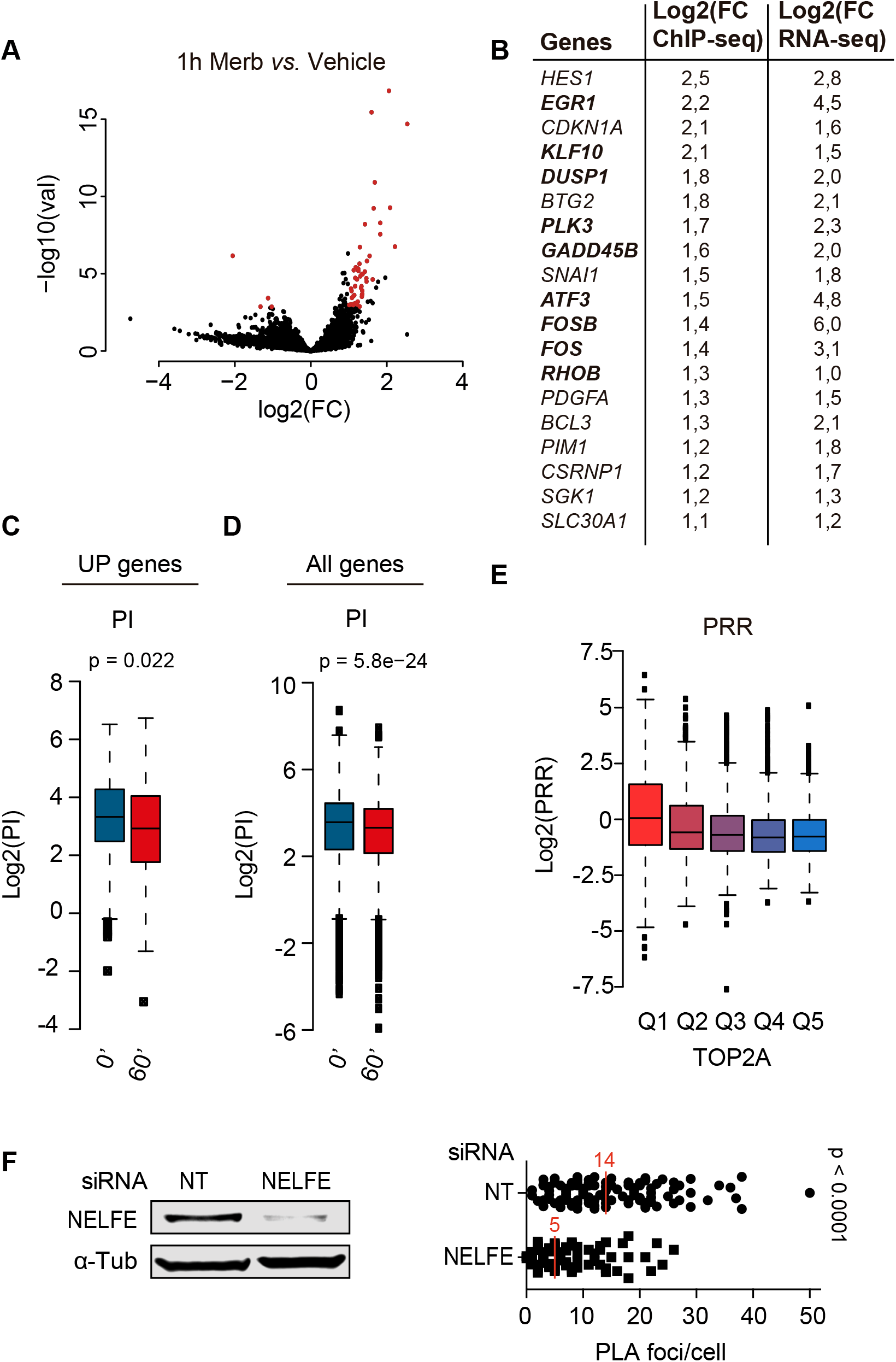
Merbarone treatment changes the distribution of Pol II. (A) Volcano plot of transcriptional changes in serum-starved RPE-1 cells upon 1 h merbarone (Merb, 200 μM) versus vehicle (DMSO) treatment, as measured by RPKM changes of Pol II ChIP-seq an the gene body (+0.5 to +2.5 kb from the TSS). The x axis represents fold-induction ratios in a log-2 scale and y axis represents p-value a log-10 scale. Genes with an absolute fold change ≥2 and an adjusted p-value ≤ 0.05 are shown in red. (B) Overlapping upregulated genes as determined by RNA seq (UP genes) and Pol II ChIP-seq. IEGs are highlighted in bold. (C and D) Pausing Index (PI) box-plot distribution for UP genes (C) and of all (10,471) transcriptionally active genes (D) following the indicated times of merbarone treatment. (E) Proximity ligation assay (PLA) quantification of TOP2A and NELFE interaction in U2OS cells transfected with non-targeting (NT) and NELFE siRNA (bottom). Westernblot validation of NELFE depletion relative to α-Tubulin loading control is shown (top). Details as in Figure 5G. Unpaired t test.

**Figure S6.**
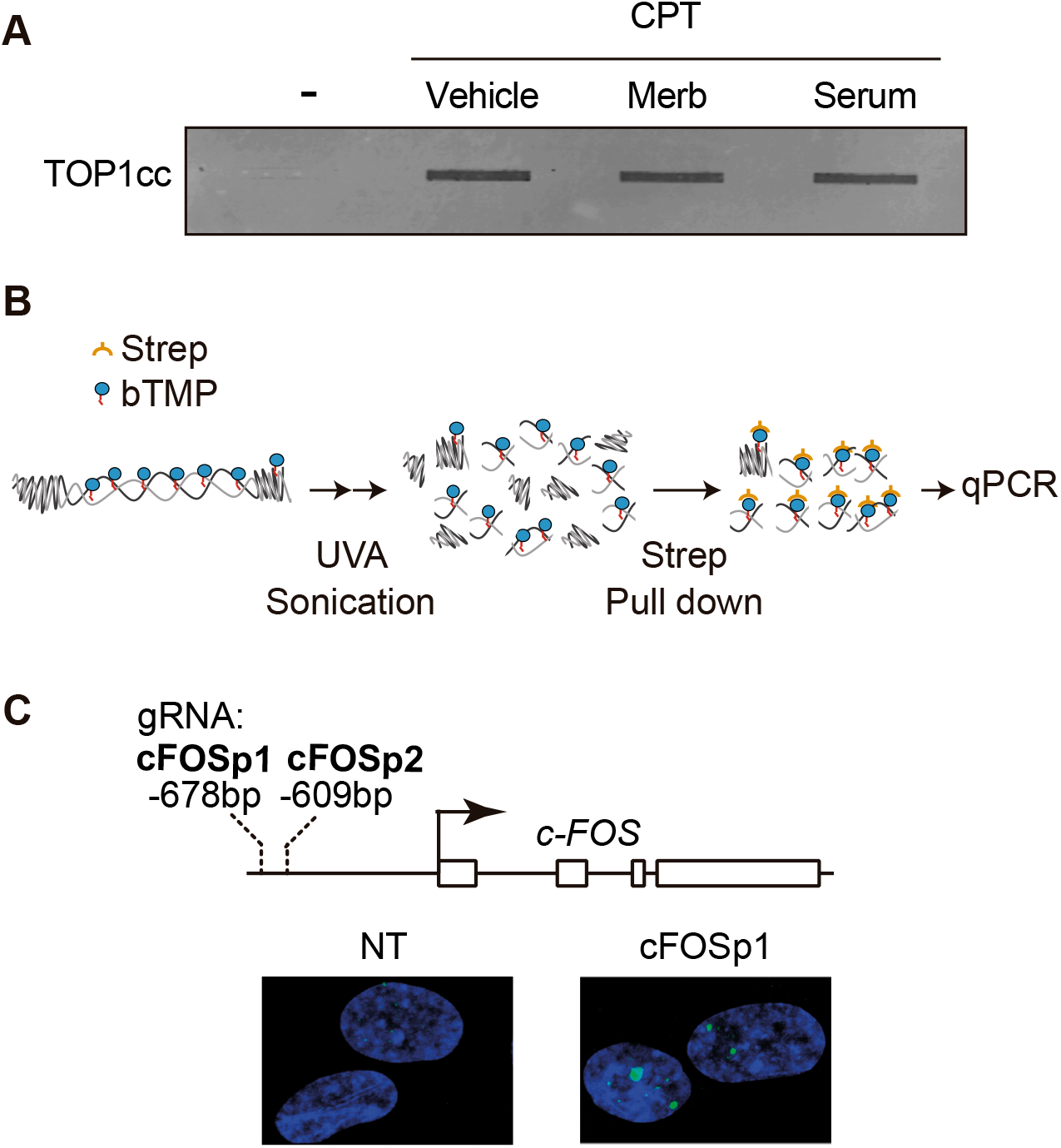
DNA supercoiling at the *cFOS* promoter. (A) Representative image (*n=3*) of TOP1 ICE assay under 30 min treatment with vehicle (DMSO), merbarone (Merb) or serum (1%) and following 5 minutes acute treatment with camptothecin (CPT, 10 μM) or vehicle (DMSO), as indicated. (B) Scheme of the experimental set-up to measure DNA supercoiling by biotin-trimethyl-psoralen (bTMP) incorporation and cross-link at the *c-FOS* gene. Following permeabilization and incubation with bTMP, cells are irradiated with UVA to crosslink psoralen to DNA. DNA is isolated, sonicated and subjected to streptavidin pull-down and qPCR amplification. (C) Diagram of the positions of the guide RNA (gRNAs) positions targeted to the *c-FOS* promoter and representative image of γH2AX (green) immunofluorescence in RPE-1 Cas9 cells 2 h after transfection with the indicated gRNA (*n = 3*). DAPI counterstaining (blue) is shown. Note that two γH2AX foci are observed in the majority of cells transfected with *c-FOS*-specific gRNA 1 (cFOSp1) and not with the non-targeting control (NT).

## STAR methods

### KEY RESOURCES TABLE

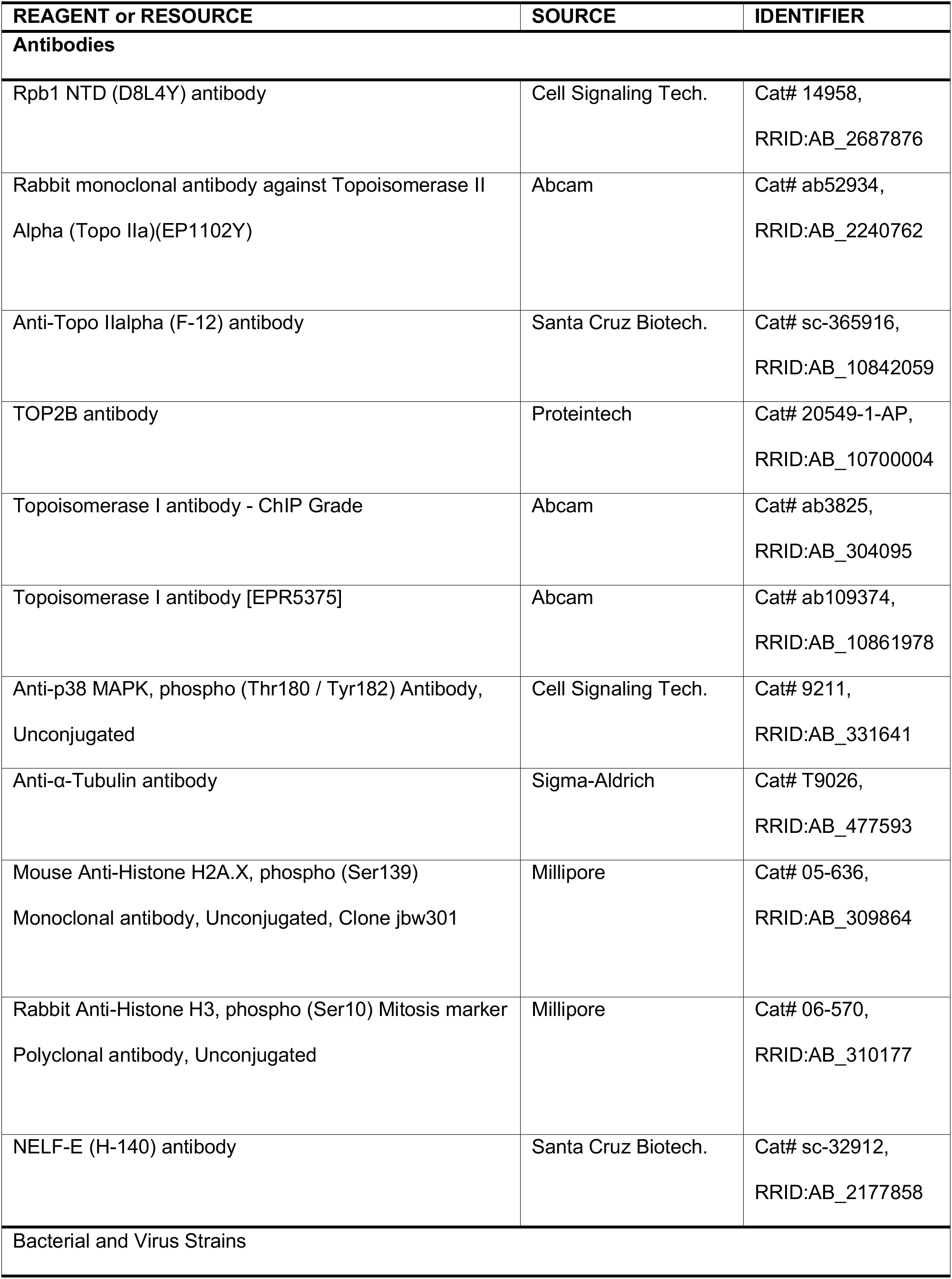
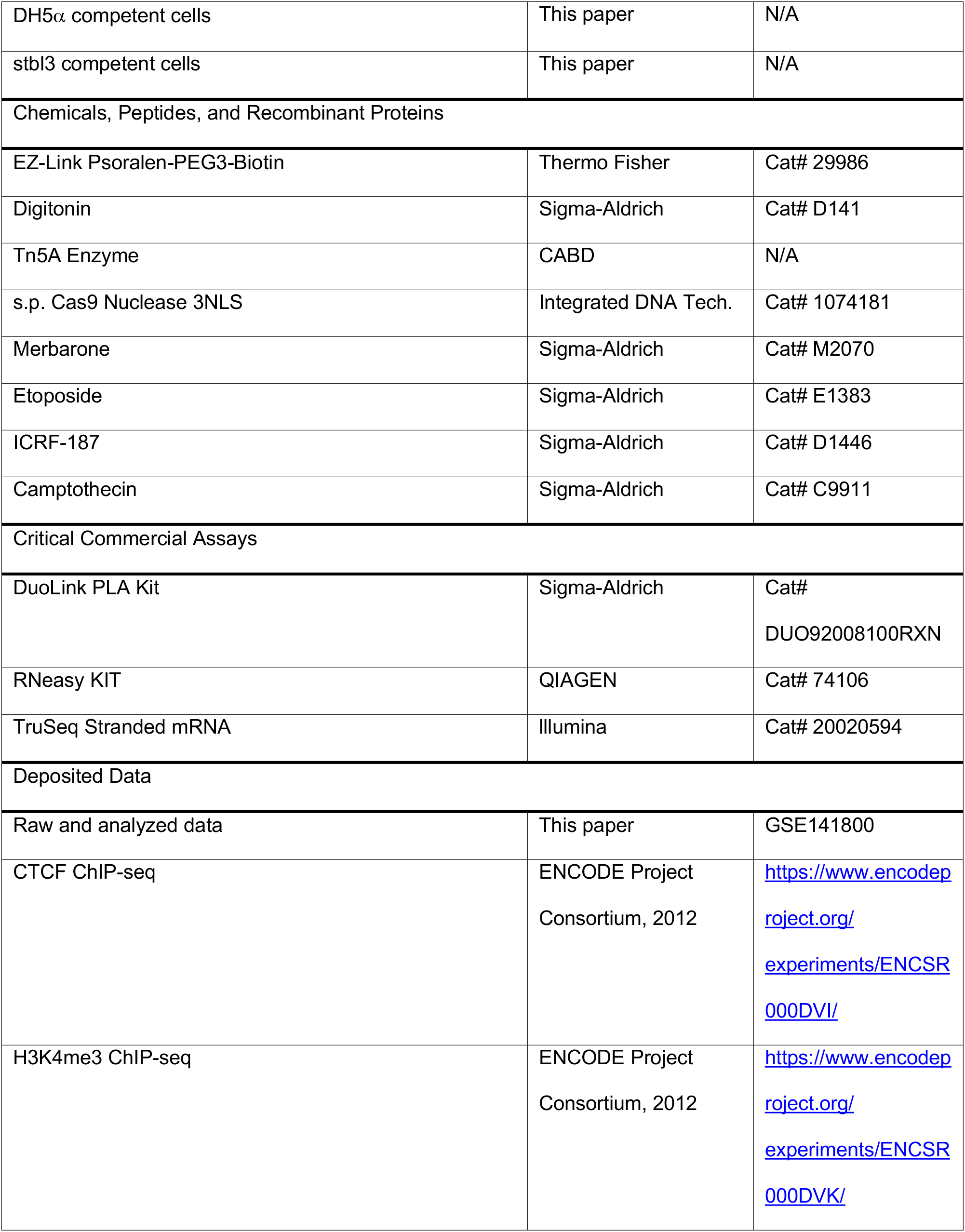
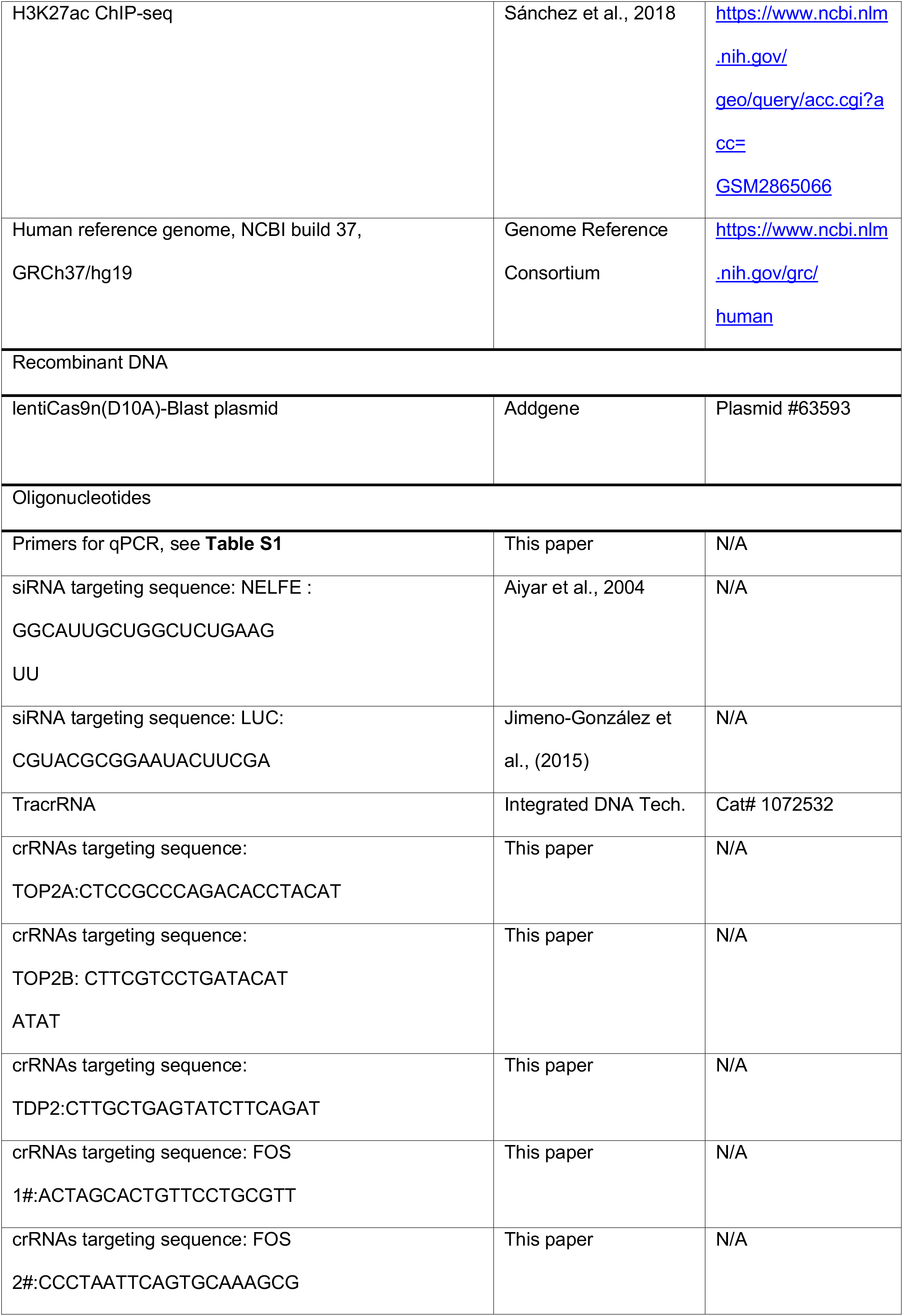
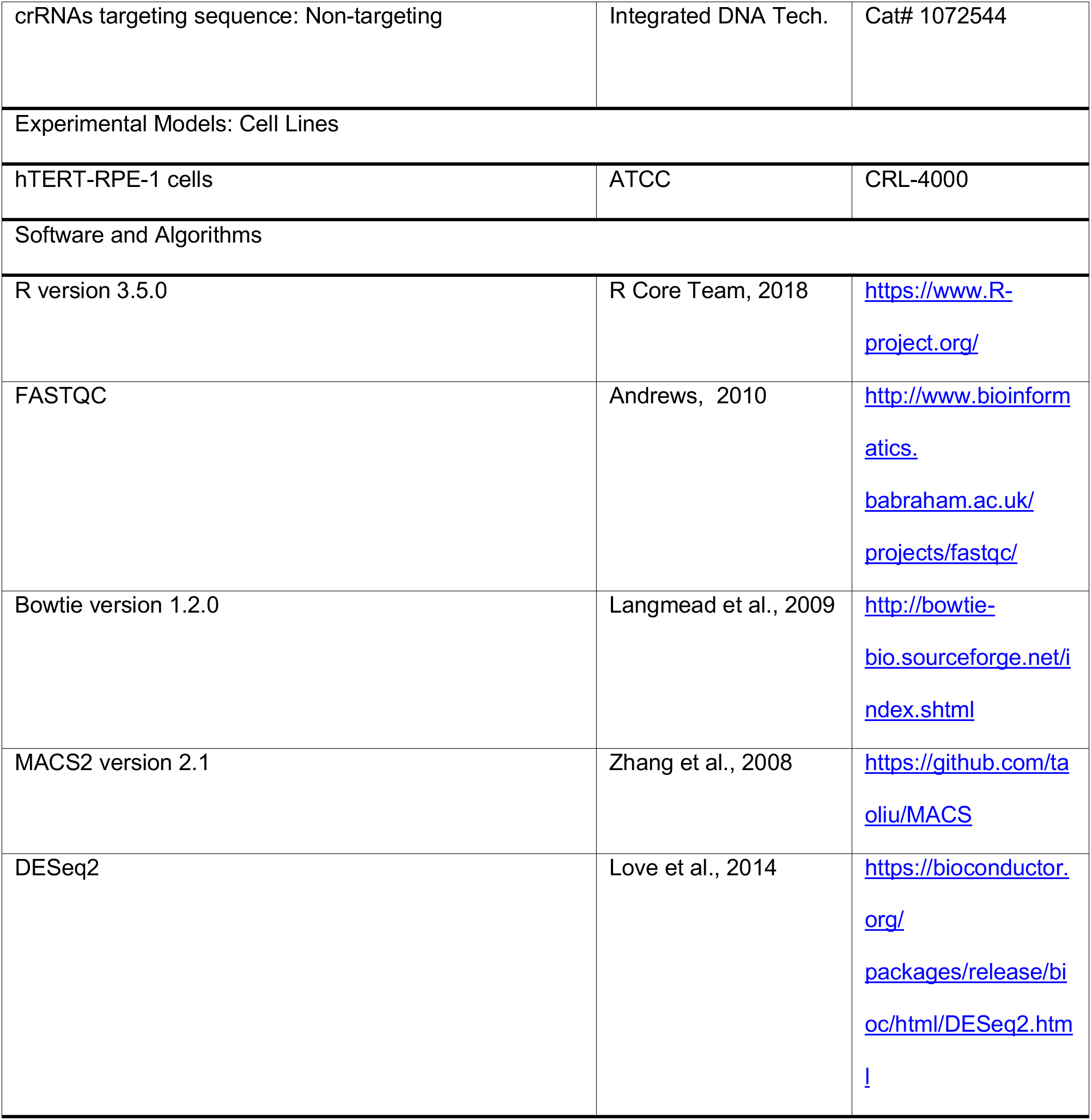

### LEAD CONTACT AND MATERIALS AVAILABILITY

Further information and requests for resources and reagents should be directed to and will be fulfilled by Lead Contact, Felipe Cortés-Ledesma.

### EXPERIMENTAL MODEL AND SUBJECT DETAILS

#### Cell lines

hTERT RPE-1 cells (ATCC), a near-diploid human cell line of female origin, were cultured in Dubelcco’s Modified Eagle’s Medium (DMEM) F-12 (Sigma) supplemented with 50 units ml-1 penicillin, 50 units ml-1 streptomycin and 10% Fetal Bovine Serum (FBS) (Sigma) at 37°C in 5% CO2 atmosphere. RPE-1 cells were serum starved by 48h incubation in the same medium, but with reduced 0.1% FBS content. Primary MEFs were isolated at day 13 p.c. and cultured in DMEM with 50 units ml-1 penicillin, 50 units ml-1 streptomycin, 15% FBS and non-essential amino acids at 37°C in 5% CO2 and 3% O2 atmosphere. HEK293T, U2OS and A549 were cultured in DMEM with 50 units ml-1 penicillin, 50 units ml-1 streptomycin and 10% FBS at 37°C in 5% CO2 atmosphere. For the generation of knockout cell lines, Cas9-overexpressing hTERT RPE-1 cell line (kindly provided by Dr. Durocher) was used. Cas9 D10A-overexpressing hTERT RPE-1 cell line was generated using lentiviral particles encoding the Cas9 D10A gene, previously produced by calcium phosphate transfection as describe (Salmon and Trono, 2006). In brief, HEK293T were transfected with a 3:2:1 mixture composed of lentiCas9n(D10A)-Blast plasmid (Addgene # 63593), p8.91 and pVSVG (Packaging plasmids), using 128 μM CaCl and 1xHBS. After 48 hours, medium was filtered through a 0.45 μm polyvinylidene difluoride (PVDF) filter (SLHV035RS, Millex-HV, Millipore). Then, viral particles were concentrated by centrifugation for 90 minutes at 22000 rpm at 4°C and stored at −80°C. The presence of mycoplasma was routinely checked with MycoAlert™ PLUS Mycoplasma Detection Kit (Lonza).

### METHOD DETAILS

#### Knock-out cell line generation

Cas9-overexpressing hTERT RPE-1 cell line was transfected with the corresponding gRNAs or non-targeting gRNA as a negative control (KEY RESOURCES TABLE),using Lipofectamine RNAiMAX (Thermo Fisher, 13778500), following the protocol provided by the manufacturer. Editing efficiency of all gRNAs was validated by in-del analysis of PCR sanger sequencing using TIDE (Brinkman et al., 2014). TOP2B^-/-^ and TOP2A^-/-^ cell lines were generated as a pool. *TDP2*^-/-^ clones were obtained by limited dilution plating in a 96-well plate.

#### Antibodies

The antibodies used are: for ChIP-seq, anti-Rpb1-NTD (Cell Signaling, D8L4Y), anti-TOP2A (Abcam, ab52934). For ICE, anti-TOP2B (Proteintech, 20549-1-AP), anti-TOP2A (Santa Cruz, SC-365916), anti-TOP1 (Abcam, ab3825). For ICE-IP, anti-TOP2A (abcam, ab52934), anti-TOP1 antibody (abcam, ab109374). For western blot analysis, anti-TOP2B (Proteintech, 20549-1-AP), anti-TOP2A (Santa Cruz, SC-365916), anti-p-p38 (Cell Signaling, 9211) and anti-tubulin (Sigma, T9026), secondary antibodies: IRDye 680-labeled anti-mouse (LI-COR Biosciences, 926-68070) and IRDye 800-labeled anti-rabbit (LI-COR BIOSCIENCES, 926-32211). For immunofluorescence, anti-γH2AX (Millipore, 05-636) and anti-H3S10p (Millipore, 06-570). For PLA, anti-NELF-E (Santa Cruz, sc32912), TOP2A (Santa Cruz, sc-365916), secondary antibodies: Alexa Fluor 488 anti-mouse (Jackson, 715-545-150), Alexa Fluor 594 anti-rabbit (Jackson, 111-585-003).

#### RNA analysis and RNA-seq

Serum-starved RPE-1 cells grown on 60mm plates were treated as required and total RNA was isolated with the RNeasy kit (Qiagen, 74106), following instructions from the manufacturer. Primers used are described in Table S1. Values were normalized to the expression of *GAPDH* housekeeping gene. For RNA-seq, total RNA (150ng) cDNA libraries were prepared using TruSeq Stranded mRNA (lllumina). Library size distribution was analyzed with Bioanalyzer DNA high-sensitive chip and Qubit. 1.4pM of each library was sequenced in NextSeq 500 HIGH-Output.

#### Western blot analysis

For protein extractions, cell pellets were resuspended in RIPA buffer (20 mM Tris-HCl (pH 7.5), 150 mM NaCl, 1% NP-40 y 1% sodium deoxycholate) supplemented with protease inhibitors and incubated on ice for 30 min with constant agitation. The lysate was then centrifuged at 14,000 rpm for 10 min at 4°C. The supernatant was sonicated using a Bioruptor (Diagenode, UCD-200) for 1 cycle of 3 minutes (high power, 30 seconds on, 30 seconds off). Protein concentration was determined by Bradford assay (Applied Biochem, A6932). 20 μg of protein was loaded home-made 10% polyacrylamide gel with SDS or 4-20% Mini-PROTEAN tris-Glycine Precast Protein Gels (Biorad, 4561096) and electroblotted onto Immobilon-FL Transfer Membranes (Millipore), after 5 minutes methanol activation. Membranes were then blocked in Odyssey Blocking Buffer (LI-COR Biosciences, 927-40000) for 1 hour and then probed with required primary antibodies for 2 hours. Membranes were washes with 0.1% Tween-20 - Odyssey Blocking Buffer and incubated with corresponding secondary antibodies (conc: 1:10.000) for 1 hour and finally, membranes were washes again with 0.1% Tween-20 - Odyssey Blocking Buffer. Membrane were analysed using Odyssey CLx and ImageStudio Odyssey CLx Software (LI-COR BIOSCIENCES, Lincoln, NE) according to the manufacturer’s protocols.

#### Chromatin Immunoprecipitation

Chromatin Immunoprecipitation were performed as previously described (Jimeno-Gonzalez et al., 2015; Subtil-Rodriguez and Reyes, 2010). Briefly, serum-starved RPE-1 cells were crosslinked with 1% formaldehyde for 10 minutes at 37°C. Crosslinking reaction was quenched with 125mM glycine for 5 minutes. Cell pellets were resuspended in 2.5 ml lysis buffer A (5 mM Pipes pH 8.0, 85 mM KCl, 0.5% NP40) supplemented with protease inhibitors and incubated for 10 minutes on ice. Chromatin was obtained by centrifugation at 4000 rpm for 5 minutes at 4°C. Nuclear fraction were resuspended in 1 ml of lysis buffer B (50 mM Tris HCl pH 8.1, 1% SDS, 10 mM EDTA, 1 mM PMSF) supplemented with protease inhibitors. Chromatin was sonicated using a Bioruptor (Diagenode, UCD-200), 10 cycles of 30” sonication (high level) and 30” of pause on icecold water. 50μl of sonicated chromatin was reverse-crosslinked using Proteinase K in PK buffer (0.5%SDS, 50mM Tris-Cl, 100mM NaCl, 1mM EDTA) at 65°C overnight. After phenol chloroform extraction, DNA fragmentation was analysed on 1.2% agarose gel. For each inmunoprecipitation, 50μg of pre-cleared chromatin and 4μg of the specific antibody was used in IP buffer (0,1% SDS, 1% TX-100, 2mM EDTA, 20 mM TrisHCl pH8, 150 mM NaCl) at 4°C o/n and then with 40 μl of pre-blocked (1 mg/ml BSA) Dynabeads protein A (ThermoFisher). Beads were sequentially washed with Wash buffer 1 (20 mM Tris HCl pH 8.1, 0.1% SDS, 1% Triton x-100, 2 mM EDTA, 150 mM NaC), Wash buffer 2 (20 mM Tris HCl pH 8.1, 0.1% SDS, 1% Triton x-100, 2 mM EDTA, 500 mM NaCl) Wash buffer 3 (10 mM Tris HCl pH 8.1, 1% NP-40, 1% NaDoc, 1 mM EDTA, 250 mM LiCl), all supplemented with protease inhibitors and twice with TE-buffer (10 mM Tris-HCL pH8, 1 mM EDTA pH8),. ChIPmentation was carried out as previously described (Schmidl et al., 2015), using Tn5A Enzyme provided by the Proteomic Service of CABD (Centro Andaluz de Biología del Desarrollo). Tagmented DNA was then eluted with 1% SDS in TE at 65°C for 10 minutes and protein was degraded with Proteinase K for 2 hours at 37 °C. DNA was purified using Qiagen PCR purification Kit (Qiagen, 28106). Libraries were amplified for N-1 cycles (being N the optimum Cq determined by qPCR reaction) using NEBNext High-Fidelity Polymerase (New England Biolabs, M0541). Libraries were purified with Sera-Mag Select Beads (GE Healthcare, 29343052) and sequenced using Illumina NextSeq 500 and single-end configuration.

#### In vivo Complex of Enzyme (ICE) and ICE-IP

DNA topoisomerase cleavage complexes were analysed as previously described (Schellenberg et al., 2017). For the induction of cleavage complexes, serum-starved RPE-1 cells were treated as required followed by 400 μM Etoposide (Sigma, E1383), 10 μM Camptothecin (CPT, C9911) (Sigma) or DMSO vehicle (Applichem, A1584) for 5 minutes. Cells were immediately lysed using 1% (w/v) N-Lauroylsarcosine sodium salt (Sigma-Aldrich, L7414) in TE buffer supplemented with protease inhibitors. After homogenization, 0.67 g/ml CsCl (Applichem-Panreac, A1098) was added and lysated were then centrifuged at 57,000 rpm for 20 h at 25 °C using 3.3 ml 13 x 33 polyallomer Optiseal tubes (Beckman Coulter) in a TLN100 rotor (Beackman Coulter).

For ICE-IP, 40 μg of ICE material was digested overnight with 0.8 U/μl Pst I (NEB, R0140). Samples were incubated at 80°C for 20min to inactivate Pst I and then diluted 1/10 in IP buffer (0.1% SDS, 1% TX-100, 2mM EDTA, 20 mM TrisHCl pH8, 150 mM NaCl). Samples were then incubated overnight at 4 °C with 4 μg of the required primary antibody and then with 40 μl of pre-blocked (1 mg/ml BSA) Dynabeads protein A (Thermo Fisher) were added and IPs were incubated for 2 hours at RT. Beads were then washed with Wash solution 1 (20 mM Tris HCl pH 8.1, 0.1% SDS, 1% Triton x-100, 2 mM EDTA, 150 mM NaCl), Wash solution 2 (20 mM Tris HCl pH 8.1, 0.1% SDS, 1% Triton x-100, 2 mM EDTA, 500 mM NaCl), Wash solution 3 (10 mM Tris HCl pH 8.1, 1% NP-40, 1% NaDoc, 1 mM EDTA, 250 mM LiCl), all supplemented with protease inhibitors, and finally with TE. DNA was then eluted with 1% SDS in TE at 65°C for 10 minutes and protein was degraded with Proteinase K for 2 hours at 37 °C. Finally, DNA was purified using Qiagen PCR purification Kit (28106, Qiagen) and analyzed by qPCR.

#### Immunofluorescence

Serum-starved RPE-1 cells grown on coverslips were fixed with 4% PFA-PBS for 10 minutes at RT. Immunofluorescence was carried out as previously described (Alvarez-Quilon et al., 2014). In brief, after permeabilization (2 minutes in PBS-0.2% Triton X-100), Cells were blocked with 5% BSA-PBS for 30 minutes and incubated with the required primary antibodies in 1% BSA-PBS for 1 hour. Cells were then washed (three times in PBS-0.1% Tween 20) and incubated with the corresponding AlexaFluor-conjugated secondary antibodies (1/1,000 dilution in 1% BSA-PBS) for 30 minutes and washed again. Finally, samples were counterstained with DAPI (Sigma, D9542) and mounted in Vectashield (Vector Labs). γH2AX foci per cell (40 cells per condition and experimental repeat) were manually counted (double-blind). pH3S10 signal was quantified with Metamorph software (100 cells per condition and experimental repeat).

#### Proximity Ligation Assay (PLA)

U2OS cells were fixed in 4% PFA for 10 min. DuoLink PLA Kit (Sigma-Aldrich, DUO92008100RXN) was used following the protocol from the manufacturer. Foci per cell (40 cells per condition and experimental repeat) were manually counted (double-blind).

#### bTMP-incorporation assay

Biotinylated-trimethylpsoralen (bTMP) incorporation was measured as previously described (Naughton et al., 2013). Briefly, serum-starved RPE-1 cells were treated as required for 10 minutes prior to the addition of 200μM EZ-Link Psoralen-PEG3-Biotin (Thermo, 29986) and 0,01% digitonin () for 5 minutes at 37°C in the dark. bTMP was cross-linked to DNA with 360 nm UV irradiation for 20 minutes on ICE. DNA was purified using Proteinase K in PK buffer (0.5%SDS, 50mM Tris-Cl, 100mM NaCl, 1mM EDTA) at 65°C overnight, followed by phenol:chloroform:isoamylalcohol extraction. After RNAse treatment and phenol:chloroform:isoamylalcohol extraction, DNA was fragmented by sonication using Bioruptor (Diagenode, UCD-200), 10 cycles of 30” sonication (high level) and 30” of pause on ice-cold water. The Biotinylated-psoralen DNA complex in TE was incubated with avidin conjugated to magnetic beads (Thermo, 6560) for 2 hours at room temperature, and then overnight at 4°C. Beads were washed sequentially for 5 minutes each at room temperature with Wash solution 1 (20 mM Tris pH 8.1, 2 mM EDTA, 150 mM NaCl, 1% Triton X-100 and 0.1% SDS), Wash solution 2 (20 mM Tris pH 8.1, 2 mM EDTA, 500 mM NaCl, 1% Triton X-100 and 0.1% SDS) and Wash solution 3 (10 mM Tris pH 8.1, 0.25 M LiCl, 1 mM EDTA, 1% NP40 and 1% deoxycholate). Beads were then washed twice with TE for 5 minutes. To extract DNA and to release psoralen adducts, samples were treated for 10 mins at 90°C in 50 μl 95% formamide - 10 mM EDTA. Samples were then brought to 200 μl with mQ water and DNA was purified using a Qiagen PCR purification Kit (28106, Qiagen) and analysed by qPCR.

#### High-throughput sequencing analysis

Sequence reads were demultiplexed, quality filtered with FastQC (Andrews, 2010) and mapped to the human genome (hg19) using Bowtie 1.2 (Langmead et al., 2009). We used option “-m 1” to retain those reads that map only once to the genome. Each individual sample contributed with the same number of reads in the ChIP-seq final merged sample. For the computation of ChIP-seq binding sites (peaks), we used MACS2 (Zhang et al., 2008) with option “-q 0.01”. We used the R package DESeq2 (Love et al., 2014) to identify differentially expressed genes from RNA-seq data following authors guidelines. Only genes with associated adjusted p-values ≤ 0.05 and absolute fold change ≥ 2 were considered as differentially expressed.

For the identification of Pol II differentially bound genes, we followed the same approach than for differentially expressed genes, this time by restricting the gene length to the region stretching 2kb from 500 bp downstream the TSS.

To estimate the level of Pol II recruitment, we used the so-called “pausing index” (PI) and “pause release ratio” (PRR). PI is defined as the ratio of Pol II enrichment within the promoter to that in the gene body, while PRR is an inverse of PI that restricts the gene body to the first 2kb downstream the TSS. For the estimation of both parameters, we based our strategies on Chen et al. (Chen et al., 2015a) and Day et al. (Day et al., 2016). To calculate PI, the level of Pol II within the promoter was computed as the sum of ChIP-seq reads in 400 bp surrounding the TSS. The level of Pol II within the gene was computed as the average number of reads in 400 bp windows throughout the gene body, from 200 bp downstream the TSS. Finally, the PI was estimated as the ratio of both measures. To calculate PRR, Pol II level within the promoter was estimated as the sum of ChIP-seq reads from 100 bp upstream to 300 bp downstream of the TSS. Gene body Pol II level was computed as the sum of reads within the region stretching from 300 bp downstream of the TSS to 2 kb downstream of the TSS. After normalizing each value by the corresponding window sizes, the PRR was estimated as the level of Pol II within the gene body divided by the level of Pol II within the promoter.

For Gene-filtering, we started from the whole set of protein-coding transcripts associated to Ensembl-annotated genes (GRCh37, release 75), from which we kept only those having a peak of Pol II (see peak calling section) overlapping with the region from 0 to 500 bp downstream of the TSS. If several transcripts of the same gene were found to match this condition, the one whose TSS was closest to the Pol II peak was selected. In order to account for potential false negatives when calling Pol II peaks, some genes with no associated Pol II peak were also selected if the window from 0 to 500 bp downstream of the TSS was found to: 1) have a value of Pol II reads per million (RPM) larger than 0.5, or 2) have a value longer than 0.5 in the difference of H3K4me3 RPM and the corresponding input. Finally, genes that were closer than 1kb of another gene or were smaller than 2kb were excluded. We ended up keeping 9,588 human genes (~ 42% of total).

Regulatory regions, namely enhancers, promoters, and insulators, were defined as follows. For promoters, the whole set of transcripts associated to Ensembl-annotated genes were considered. Then, promoters were defined as ± 3 kb from the TSSs. Enhancers were defined as H3K27ac peaks not overlapping with a promoter, and insulators as CTCF peaks not overlapping with promoters and enhancers.

ChIP-seq averaged reads around TSSs were computed using the R package *bamsignals* (Mammana, 2019) and smoothed using a Gaussian smoothing kernel with the R function *ksmooth*, respectively. To generate the profile of TOP2A ChIP-seq signal around randomized genes, the 148 up-regulated genes upon 2h of merbarone treatment were first considered. Then, 50 sets of 148 genes were randomly selected from the human genome and TOP2A ChIP-seq reads were counted around the TSSs. Finally, the median of such read counts was smoothed and plotted.

Publicly available sequencing data used in this study include ChIP-seq of several proteins and post-translational modifications: CTCF (ENCSR000DVI), H3K4me3 (ENCSR000DVK) and H3K27ac (Sánchez et al., 2018). CTCF and H3K4me3 BAM files (hg19) were batch-downloaded from ENCODE. H3K27ac and H3K27me3 raw sequencing files were processed as described above (Langmead et al., 2009).

The R package TopGO (Alexa, 2019) was used to calculate the significance of Gene Ontology (GO) terms associated to differentially expressed genes after merbarone treatment. We computed enrichments using the Fisher’s exact test and the default algorithm (weight01), which is a hybrid between the ‘elim’ and the ‘weight’ algorithms described (Alexa et al., 2006). To perform hypergeometric-based tests, there is a need for defining a ‘gene universe’ (which can be conceptualized as the number of balls in an urn) and a set of ‘interesting genes’ from that universe. To define the gene universe, we started from expressed genes, which were defined as genes for which the sum of RNA-seq reads (combining the replicates) overlapping exons was larger than 10. Then, the gene universe was defined as expressed genes mapping to at least one GO term and the set of interesting genes as differentially expressed genes upon merbarone treatment.

#### Data and materials availability

Data are available in the main text, supplementary materials and auxiliary files. Nextgeneration sequencing data have been deposited at Gene Expression Omnibus with accession numbers GSE141799 and GSE141800.

## Supplementary Materials

Figures S1-S6

Table S1

Data S1

